# Intrinsically disordered proteins SAID1/2 condensate on SERRATE/ARS2 for dual inhibition of miRNA biogenesis in Arabidopsis

**DOI:** 10.1101/2022.09.13.507836

**Authors:** Baoshuan Shang, Lin Wang, Xingxing Yan, Yanjun Li, Changhao Li, Chaohua Wu, Tian Wang, Xiang-Guo, Sukwon Choi, Tianru Zhang, Ziying Wang, Chun-Yip Tong, Taerin Oh, Xiao-Zhang, Zhiye Wang, Xu Peng, Xiuren Zhang

**Author notes:** Equal contribution (Baoshuan Shang, Lin Wang, Xingxing Yan).

## Abstract

Intrinsically disordered proteins (IDPs) SAID1/2 are hypothetic dentin sialophosphoprotein-like proteins, but their true functions are unknown. Here, we identified SAID1/2 as negative regulators of SERRATE(SE)/ARS2, a core factor in miRNA biogenesis complex (microprocessor). Loss-of-function double mutants of *said1; said2* caused pleiotropic developmental defects and thousands of differentially-expressed genes that partially overlapped with those in *se. said1; said2* also displayed increased assembly of microprocessor and elevated accumulation of miRNAs. Mechanistically, SAID1/2 promotes PRP4KA-mediated phosphorylation of SE, leading to its degradation in vivo. Unexpectedly, SAID1/2 have strong binding affinity to hairpin-structured pri-miRNAs and can sequester them from SE. Moreover, SAID1/2 directly inhibit pri-miRNA processing by microprocessor in vitro. Whereas SAID1/2 did not impact SE subcellular compartmentation, the proteins themselves exhibited liquid-liquid phase condensation that is nucleated on SE. Thus, we proposed that SAID1/2 reduce miRNA production through hijacking pri-miRNAs to prevent microprocessor activity while promoting SE phosphorylation and its destabilization in Arabidopsis.

## Introduction

Up to date, 5386 of 27619 protein-coding genes have not been assigned any function and biological process in Arabidopsis (https://www.arabidopsis.org). Among the genes are 11 dentin sialophosphoprotein (DSPP) and DSPP-like (DSPPL) proteins annotated. In mammals, DSPP is a precursor protein that is broken down into several components to become the dentin extracellular matrix of the tooth (Yamamoto et al., 2015). DSPPs are also considered as small integrin-binding ligand N-linked glycoproteins, and exhibit correlations with aggressiveness in human cancers (Bellahcene et al., 2008). Whether DSPPs are secreted into extracellular matrix to strengthen plant issues, or whether the proteins are engaged in other functions are unknown.

microRNAs (miRNAs), a large family of small non-coding regulatory RNAs, play critical roles in numerous biological processes, including development, defense, and adaptation to changing environment (Li et al., 2017; Song et al., 2019). miRNAs function through Argonaute (AGO)-containing RNA induced silencing complexes (RISCs). By using miRNAs as guides, AGO proteins specify the sequence-complementary target mRNAs to repress their expression (Li et al., 2013; Ma et al., 2018a). In animals, miRNAs are produced through the sequential activity of microprocessor in the nucleus and Dicer complex in the cytoplasm (Jin et al., 2020; Kwon et al., 2016; Liu et al., 2018; Partin et al., 2020). In plants, miRNAs are initially processed from primary substrates (pri-miRNAs) to generate miRNA precursors (pre-miRNAs), which are furthermore cut into miRNA/miRNA* for incorporation into AGO proteins (Moro et al., 2018). The entire biogenesis processes are fulfilled by microprocessor/dicing complex that is minimally comprised of DCL1 and a dsRNA-binding protein, HYL1 (Zhu et al., 2013).

Since precise processing and accumulation homeostasis of miRNAs are critical for their proper functionality in biology, biogenesis and metabolism of miRNAs are fine-tuned through multiple regulatory layers. Multiple genetic pathways to control miRNA homeostasis converge on the multifunctional protein SE in Arabidopsis. SE has been best known to partner with DCL1 and HYL1 to produce miRNAs (Grigg et al., 2005; Yang et al., 2006). Earlier reports showed that SE could directly promote the enzymatic activity and accuracy of DCL1 in microprocessor reconstitution assay in vitro (Dong et al., 2008; Iwata et al., 2013). Recent studies propose that SE might act as a scaffold or mediate liquid-liquid phase separation to recruit the core processing machinery including DCL1/HYL1 to the proper RNA substrates, or *vice versa*, to generate miRNAs *in vivo* (Machida et al., 2011; Xie et al., 2021; Yang et al., 2010; Zhu et al., 2013). Moreover, the formation of SE-scaffolded dicing bodies in the nucleoplasm can be regulated by TREX-2 among other cellular factors (Zhang et al., 2020). On the other hand, SE can also fulfill negative regulation to miRNA production. For instance, SE recruits SWI2/SNF2 ATPase subunit CHR2 to remodel secondary structure of pri-miRNAs to inhibit miRNA production (Wang et al., 2018). SE also interacts with the nuclear exosome targeting (NEXT) complex to degrade pri-miRNAs to fine tune miRNA production in Arabidopsis (Bajczyk et al., 2020). Of note, SE-mediated decay of pri-miRNAs via the exoribonuclease XRN2/XRN3 can be reversed by Modifier of SNC4-Associated Complex (MAC5) (Li et al., 2020a). Similarly, the mammalian ortholog of SE, arsenic resistance protein 2 (Ars2) also participate in RNA silencing (Gruber et al., 2009; Sabin et al., 2009). Besides RNA silencing, SE/Ars2 also contributes to other aspects of RNA metabolism, for instance pre-mRNA splicing, biogenesis of non-coding RNAs, RNA transport and RNA stability (Gruber et al., 2012; Hallais et al., 2013; Laubinger et al., 2008 et al., 2010; Machitani et al., 2020; Melko et al., 2020; Raczynska et al., 2014; Thillainadesan, 2020). In addition, SE/Ars2 protein acts as a transcriptional factor, regulating expression of transposons (Ma et al., 2018b), and protein-coding genes (Andreu-Agullo et al., 2012; Speth et al., 2018; Yin et al., 2020; Yu et al., 2020).

Given the critical roles of SE/Ars2 in RNA metabolism, regulation of the proteins themselves becomes pivotal to secure the proper accumulation and processing of the target transcripts. We have recently found that SE/Ars2 are IDPs and perform their functions through various macromolecular complexes. Unpacked or excessive SE/Ars2 are scavenged by 20S Proteasome Alpha Subunit G1 (PAG1) or possibly PSMA3, the mammalian ortholog of PAG1, which are key components of 20S core proteasome, for degradation via a ubiquitin-independent pathway (Fedorova et al., 2011; Li et al., 2020c). We further demonstrated that pre-mRNA processing 4 kinase A (PRP4KA) phosphorylates SE (Wang et al., 2022). Once hyper-phosphorylated, SE has reduced binding affinity to HYL1 and is readily degraded by 20S proteasome. Phosphorylation of SE via PRP4KA can quickly clear accumulated SE to secure its proper amount. Thus, RNA metabolism can be regulated through controlling homeostasis of the key microprocessor component accumulation in eukaryotes at a posttranslational level (Wang et al., 2022).

Here, we reported two IDPs, DSPPL9 and DSPPL11, as novel SE partners to repress miRNA biogenesis. To better reflect the biological roles of the two proteins in plants, we renamed them as SAID1 (SE-Associated Inhibitors with Dwarfism appearance in the mutants) and SAID2. While single *said1* and/or *said2* T-DNA and CRISPR-Cas9 knock-out lines displayed subtle morphological defects, *said1; said2* double mutants exhibited twisted leaves and short statue but with seemly normal seed setting. Different from *se*, the double mutant generally contained elevated miRNA accumulation and reduced miRNA target expression compared with Col-0. We further found that SAID1/2 triggered phosphorylation and degradation of SE in vivo, resulting in the reduced assembly of microprocessor. Moreover, SAID1/2 had stronger binding affinities to pri-miRNAs and were able to sequester pri-miRNAs from SE. In addition, SAID1/2 could also directly inhibit the activity of the microprocessor in pri-miRNA processing in vitro. For in vivo, the two proteins displayed liquid-liquid phase separation (LLPS) condensation that can be dynamically regulated by SE protein. Thus, this study revealed novel and unexpected roles of DSPPL/SAID proteins in RNA metabolism and a new regulatory layer of miRNA biogenesis in plants, contributing to deciphering the functions of poorly understood gene family in eukaryotes.

## Results

### SAID1/2 are new *bona fide* partners of SE protein

We previously performed Mass Spectrometry (IP-MS) analysis of SE complexes and identified several novel partners that regulate SE functions and homeostasis (Li et al., 2020c; Ma et al., 2018b; Wang et al., 2022; Wang et al., 2018). Among the rest candidates of SE interactors is DSPPL9/SAID1, which is annotated as a homolog of human Dentin Sialophosphoprotein with unknown function (**Figure S1A**). This protein was also recovered in an independent IP-MS/MS of SE complexes by others (Bajczyk et al., 2020), validating the plausibility of SAID1 as a genuine partner of SE. Phylogenetic analysis suggested SAID1 has a paralog, SAID2, although they only share 15.6% identity and 23.6% similarity (**Figures S1B and S1C**). To validate the interaction of SE and SAID1/2, we performed bimolecular fluorescence complementation (BiFC) assay. Co-transfection of either nYFP-SAID1 or nYFP-SAID2 with cYFP-SE, but not with cYFP, displayed clear condensates in nuclei, reminiscent of the nuclear punctate foci formed by the positive control of nYFP-DCL1- and cYFP-SE (**Figure 1A**). Next, we validated the interaction of SE and SAID1/2 through yeast-two-hybrid (Y2H) assay (**Figure 1B**). Of note, co-transfection of BD-SE with AD-SAID1 could readily result in the growth of heathy colonies on the quadruple-deficient medium, and so could the combination of BD-SE/AD-SAID2 but with fewer colonies. This result implied that SAID2 might have less binding affinity with SE or might be less stable in vivo. Further Y2H assays with a serial of truncated variants revealed that SAID1 interacts with the C-terminal region (469-720 aa) of SE covering zinc finger domain and C-terminal unstructured tail (**Figures 1E and 1F**). Interestingly, this region is also the site for SE interaction with DCL1 and PRP4KA among other partners (Wang et al., 2022; Wang et al., 2018). On the other hand, SE interacts with the middle part (377-580aa) of SAID1 (**Figures 1C and 1D**). Of note, further fine mapping assays delineated the domain (377-407aa) of SAID1 being crucial for its interaction with SE (**Figures S1D-S1F**), implying that this region harbors certain critical amino acids or motif pivotal for the protein-protein interaction. There is low sequence homology between SAID1 and SAID2 in this region that might explain the relative reduced binding of SE by SAID2 vs SAID1 in Y2H assays.

**Figure 1.**
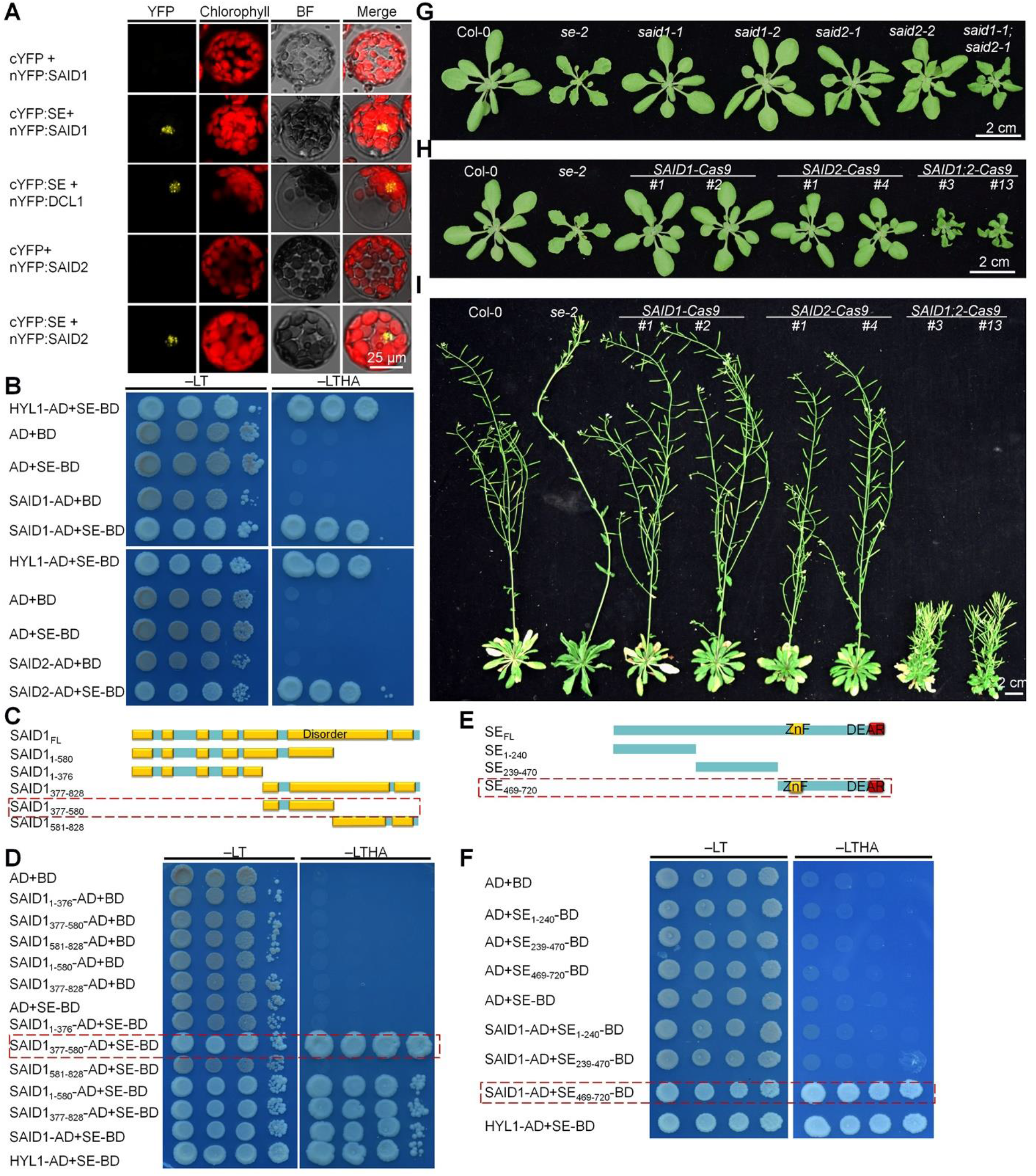
Depletion of the SE-bound partners SAID1/2 leads to pleiotropic developmental defects in plants. (**A-F**) Experimental validation of SAID1/2 as *bona fide* partners of SE by BiFC (**A**) and Y2H (**B-F**). In (**A**), various combinations of N-terminal or C-terminal YFP (nYFP, or cYFP)-tagged proteins were co-expressed in Arabidopsis mesophyll protoplasts and YFP signal indicative of the protein-protein interaction was detected. The combinations with nYFP, or cYFP alone served as negative controls. Scale bar=25 µm. At least 30 protoplasts were detected and showed similar interaction patterns. In (**B**), BD-SE and AD-SAID1/2 were co-transfected to yeast and plated on double dropout or quadruple dropout medium SD/–Leu/–Trp (−LT) or SD/– Ade/–His/–Leu/–Trp (−LTHA), respectively. In (**C-F**), Y2H assays with a serial of truncated variants of SAID1 and SE that interacts with each other. Serial truncated variants of SAID1 and SE are shown in (**C**) and (**E**), respectively, Full length (FL) and truncated forms (lower-case numbers) are shown. ZnF: zinc finger domain, GAPE: motif enriched with Gly, Ala, Pro, and Glu. In all Y2H assays, three biological repeats were performed, and representative results are shown. (**G-I**) Morphological phenotypes of *said1* and *said2* mutants. Photos were taken of 3-week-old T-DNA insertion lines (**G)** and CRISPR-Cas9 lines (**H**) and two-month-old CRISPR-Cas9 null alleles (**I**). Scale bars=2 cm.

### Inter-dependent and -independent regulatory roles of SAID1/2 and SE in gene expression

To investigate functional relevance of SAID1/2-SE interaction, we screened two T-DNA homozygote mutants for each gene (*said1-1* and *said1-2* for SALK_080762C and SALK_133343; and *said2-1* and *said2-2* for SALK_062434 and SAIL_88_D02), respectively (**Figure S2A**). The *said1-1* and *said1-2* lines did not show obvious morphological change, while *said2-1* and *said2-2* displayed mild down-curved leaves and slightly reduced stature phenotype (**Figures 1G, S2B, and S2C**). Lack of developmental defect in *said1-1* and *said1-2* might result from the fact that T-DNA insertion sites are at the last exon of the gene, and the lines might still encode truncated but partially functional protein. Alternatively, there might be the functional redundancy between SAID1 and its paralog SAID2. Indeed, the seedlings of *said1-1*; *said2-1* double mutant showed pleiotropic phenotype such as down-curved leaves, small stature, and their mature plants exhibited dwarfism and mild clustered siliques. Moreover, the developmental defects were more obvious in a long-day growth condition (**Figures S2B and S2C**). We then generated CRISPR/Cas9 knockout lines for *SAID1* and/or *SAID2* (Yan et al., 2015), with the guide RNAs targeting the first and third exons of *SAID1* and *SAID2*, respectively (**Figures S2D and S2E**). Like T-DNA insertion mutants, the single CRISPR/Cas9 knockout lines displayed normal morphology or subtle growth defect (**Figures 1H, 1I, S2F, and S2G**).

Intriguingly, most of the double knock-out lines showed severer pleiotropic phenotypes than *said1-1*; *said2-1*, namely down-curved leaves, dwarfisms, often bushy, and clustered siliques of short internode and altered phyllotaxis (**Figures 1H and 1I**). We selected two representative lines (*SAID1*; *SAID2-Cas9 #3* and *#13*) that were genotyping-validated (**Figures S2H-S2K**) and crossed them with Col-0 to clean the background for further functional analysis. Of note, computational analysis ranked 4 and 3 top off-target candidates for CRISPR-Cas9 knockout lines; however, this concern was cleared as Sanger sequencing did not recover any polymorphisms in the potential off-target sites (**Figures S2L and S2M**).

We then conducted RNA-seq of 3-week-old *SAID1*; *SAID2-Cas9 #3* (thereafter renamed as *said1-3; said2-3*) plants with *se-2* and *chr2-1* references (**Figure S3A**). Bioinformatic analysis recovered 5520 differentially expressed genes (DEGs) in *said1-3; said2-3* vs Col-0. By contrast, only 1801 DEGs were detected in *se-2* (**Figure 2A, Table S3**). This number was much lower than that in *se-2* observed in our earlier RNA-seq (Li et al., 2020c), likely due to different growing conditions, with further suggestion that the numbers of DEGs recovered in this RNA-seq might be underestimated.

**Figure 2.**
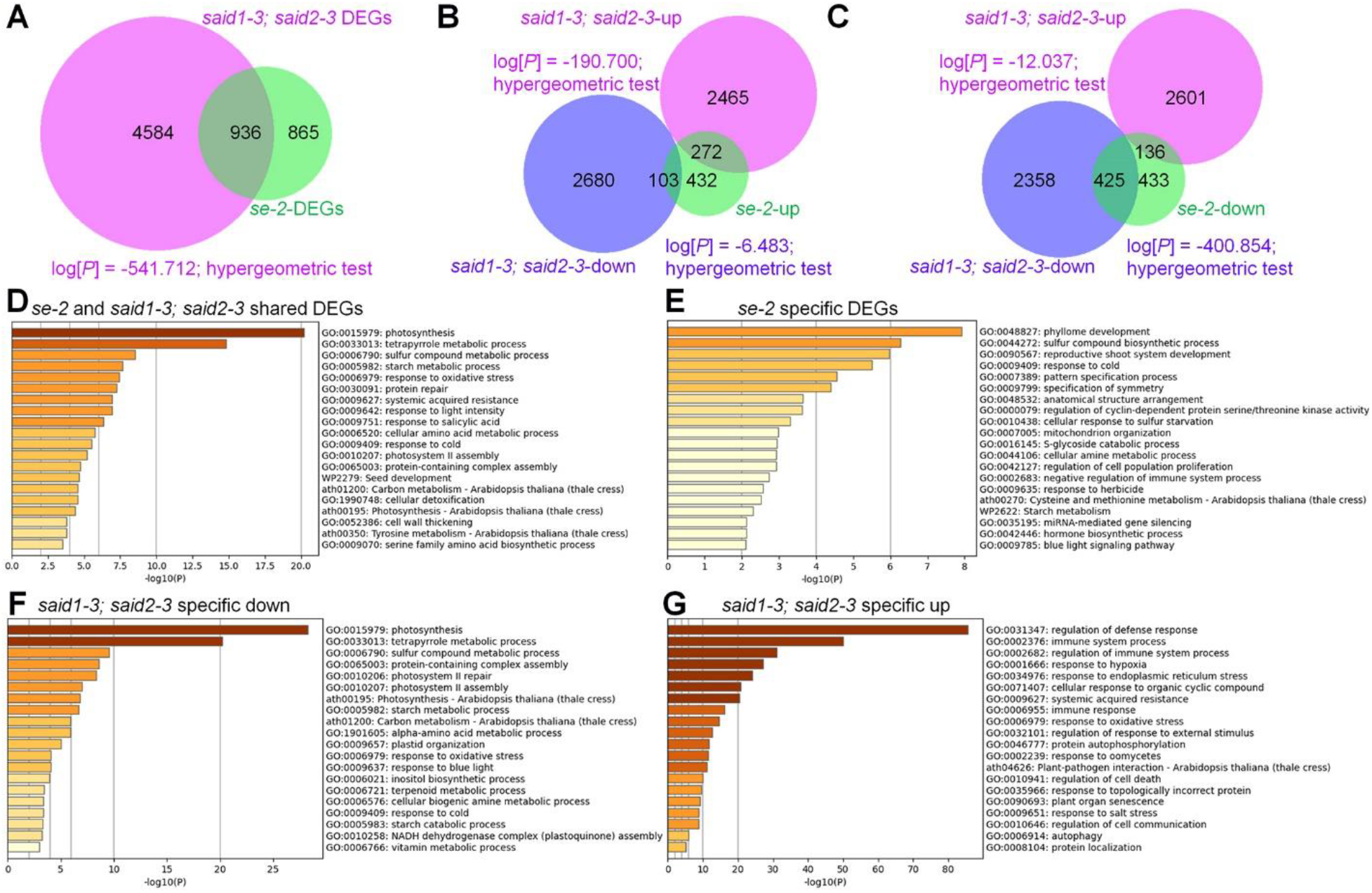
Overlapping and non-overlapping DEGs between *said1-3; said2-3* and *se-2* mutants. (**A**) RNA-seq revealed genome-wide overlapped and distinct DEGs between *said1-3; said2-3* and *se-2* mutants. (**B** and **C**) Overlapping of up-regulated and down-regulated genes between *said1; said2* and *se-2* mutants. Correlation of overlapped DEGs were tested by hypergeometric test. For (**A-C**), FDR< 0.05 as cut-off. n=three biological replicates. (**D-G**) GO enrichment of the DEGs that are shared or specific for *said1-3; said2-3* and *se-2*, respectively.

Among 5520 DEGs, 2737 and 2783 genes were either up-regulated or down-regulated in *said1-3; said2-3* vs Col-0, respectively (**Figures 2A-2C**). Notably, 936 of 5520 DEGs in *said1-3; said2-3* were shared with the ones observed in *se-2*. Among the shared DEGs, they appeared to exhibit more concordance than difference between *se-2* and *said1-3; said2-3* mutants. For instance, 272 genes, representing ~34% of 807 upregulated ones in *se-2*, were accordantly upregulated in *said1-3; said2-3*, whereas only 103 genes, representing 13% of 807 upregulated genes in *se-2*, were downregulated in *said1-3; said2-3* (**Figure 2B**). On the other hand, 425 of 994 downregulated in *se-2* (~43%) were repressed in *said1-3; said2-3* and only up to 14% of *se-2*-repressed genes were upregulated in the double mutant (**Figure 2C**). These results suggested that *SAID1/2* and *SE* might have shared targets. Although more than 50% of DEGs in *se-2* overlapped with those of *said1-3; said2-3*, a majority of DEGs in *said1-3; said2-3* were not observed in *se-2* (**Figures 2A-2C**). In addition, several SE-dependent alternative splicing genes did not show obvious defect of pre-mRNA splicing in *said1-3; said2-3* (**Figure S3B**). These results suggested that *SAID1/2* and *SE* might also have inter-independent regulatory roles in gene expression in vivo.

The DEGs shared between *said1-3; said2-3* and *se-2* were mostly enriched in the pathways of photosynthesis, metabolism of carbon and starch, and stress responses (**Figure 2D**). This pattern is in lines with the low biomass production of both mutants (**Figures 1G-1I**). Gene Ontology (GO) analysis of the genes specifically downregulated in *said1-3; said2-3* pointed out that the most enrichment of DEGs were related to metabolic and cellular processes and response to stimulus, followed by regulation of rhythmic and other biological processes, whereas DEGs in *se-2* are related to phyllome development and organ polarity among others (**Figures 2E and 2F**). In consistent to this regulation is predicted interaction of SAID1 with the oscillator TIC (Time for Coffee) that regulates circadian rhythm and the potential interaction of SAID2 interaction with CSP3 (Cold shock domain-containing protein 3) in plants (https://string-db.org). Unexpectedly, the significantly overrepresented genes in *said1-3; said2-3* were engaged in immune responses, systemic acquired resistance, and plant responses to various hormones or environmental stresses (**Figure 2G**). The fact that the defense- and other stress-responsive mRNAs accumulate despite growth in a normal physiological condition with simultaneous repression of mRNAs required for general biological processes in *said1-3; said2-3* vs Col-0 implied that, wild-type SAID1/2, reminiscent of DHH1/DDX6-like RNA helicases (Chantarachot et al., 2020), suppress stress-responsive mRNAs under non-stress conditions to maintain growth/defense balance in plants. The DEG patten also explained the dwarfism phenotype characteristic of the double mutants.

### SAID1/2 repress miRNA accumulation *in vivo*

We next conducted small RNA-seq (sRNA-seq) to evaluate miRNA profiles in *said1* and/or *said2* mutants (**Figure S4A**). Whereas most of annotated miRNA species were down-regulated in *se-2*, small proportions of miRNAs were differentially accumulated in *said1-1* (**Figures 3A, and S4B**). Of note, ~80 miRNAs were up-regulated and ~30 miRNAs down-regulated in *said2-1*, and the relatively increased differentially accumulated number of miRNAs is consistent with the mild morphological change in *said2-1* relative to *said1-1* (**Figures 1G, S2B, S2C, and S4C**). Importantly, the *said1-3; said2-3* line showed ~110 up-regulated miRNAs, with less than 50 of down-regulated miRNAs (**Figure 3B**). An independent line (*said1-13; said2-13*) also showed a similar pattern, but to a lesser extent (**Figure S4D**). Furthermore, the up-regulated miRNAs typically belong to SE-controlled canonical species (**Figure 3C**), indicating negative role of SAID1/2 to regulate miRNA accumulation. We randomly selected a few canonical miRNAs and their star strands for sRNA blot and stem-loop qRT-PCR assays. Clearly, the assays readily validated the sRNA-seq results above (**Figures 3D and 3E**). Taken together, we concluded that the two novel proteins SAID1/2 generally repress miRNA accumulation in vivo. The fact that some miRNAs are reduced in *said1-3; said2-3* implies that SAID proteins might have some regulatory roles in addition to impacting SE function.

**Figure 3.**
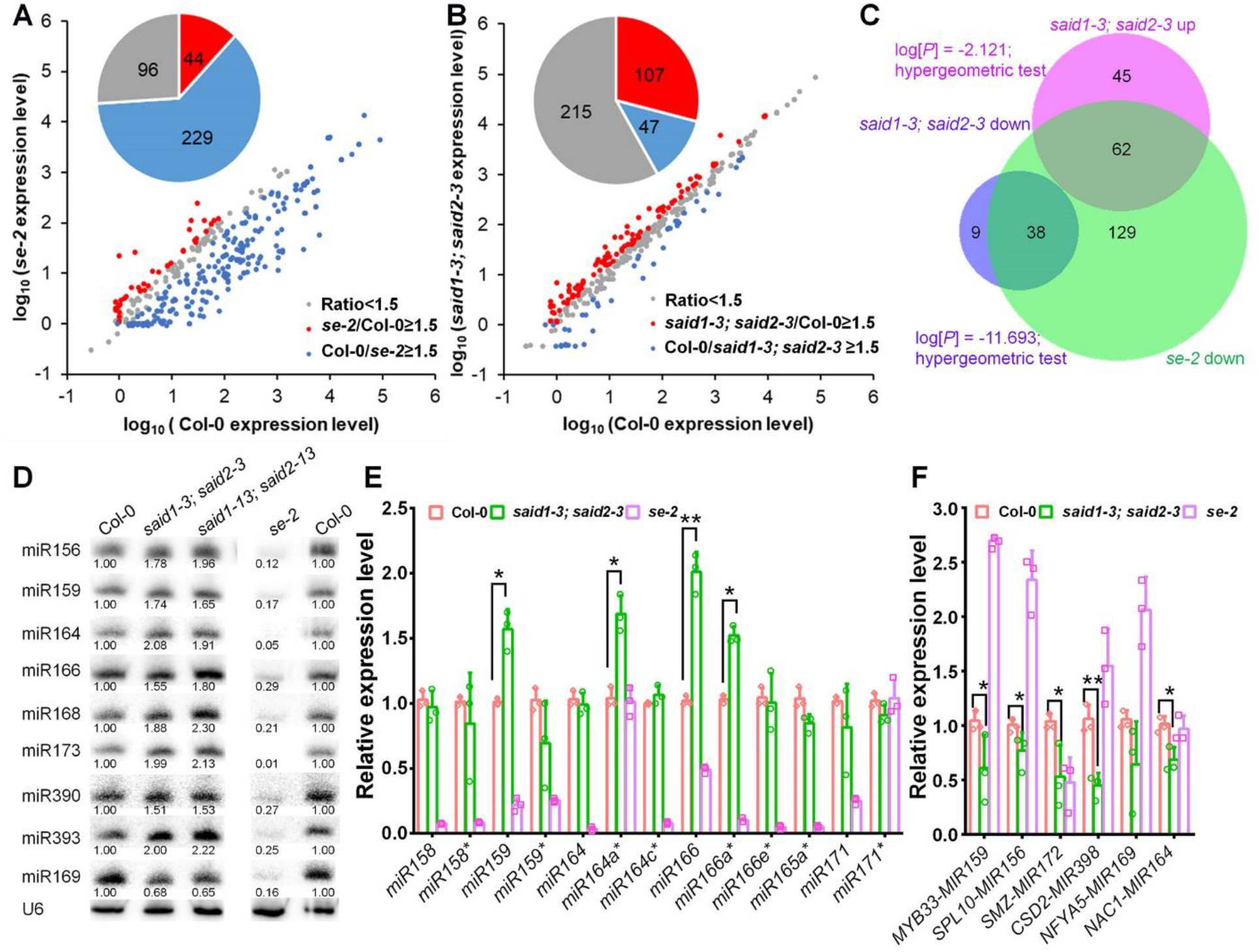
SAID1/2 negatively regulate miRNA accumulation. (**A-B**) sRNA sequencing analysis of miRNA profiling in Cas9 null mutants of *said1-3; said2-3, se-2* relative to Col-0. miRNAs with expression of at least 1.5-fold higher (mutants/Col-0 ≥ 1.5), lower (Col-0/mutant ≥ 1.5), or within <1.5-fold are indicated by red, blue, and gray dots, respectively, with the means from three biological replicates (n=3, false discovery rate (FDR) < 0.05). (**C**) Overlapping of up- and down-regulated miRNAs in *said1; said2* with SE-dependent miRNAs. Correlation of overlapped de-regulated miRNAs were tested by hypergeometric test. (**D, E**) sRNA blot (**D**) and stem-loop qRT-PCR (**E**) of selected miRNAs in indicated mutants. U6 served as loading or normalization controls in (**D**) and (**E**), respectively. The numbers below the gels (**D**) indicate the relative mean signals of miRNAs that were normalized to that of Col-0 where the value was arbitrarily assigned a value of 1. (**F**) Heat map and hierarchical clustering analysis shows that a subset of miRNA targets were similarly downregulated in *said1; said2* and *chr2-1* vs Col-0. (**G**) qRT-PCR assays validated the reduced accumulation of selected miRNA targets in *said1-3; said2-3* compared to Col-0. Transcript levels in the indicated mutants were first normalized to the internal control *EF-1α*, and then to that of Col-0. In (**E** and **G**), Data are means ± SD of three biological replicates with statistics significance determined by Student’s *t* test. *P < 0.05, **P < 0.01.

We also assessed the expression of canonical miRNA targets in the mutants by qRT-PCR assays. Indeed, selected miRNA targets displayed significantly reduced expression in *said1-3; said2-3*, but only mildly in the single mutants *said1-1* and *said2-1* (**Figures 3E, S4E, and S4F**). These results were also consistent with increased accumulation of miRNAs in the double mutants. Of note, the expression of certain miRNA targets in *said1-3; said2-3* was reduced, suggestive of elevated miRNA accumulation (**Figure 3F**), despite the observation that there was no or subtle difference in the expression of related miRNAs in the sRNA blot (or sRNA-seq) analysis (**Figures 3B and 3D**). This contrast might be caused by low sensitivity of RNA blot assays or barcode bias in the sRNA-seq (Raabe et al., 2014; Sorefan et al., 2012). We also observed that numerous miRNA targets displayed increased or relatively stable expression, indicating that these miRNA targets might be subjective to additional layers of regulation in vivo. Importantly, the overall expression pattern of miRNA targets in *said1-3; said2-3* very much resembles that in *chr2-1* (Wang et al., 2018), further suggesting that SAID1/2, like CHR2, are novel negative regulators in the miRNA pathway in vivo.

### SAID1/2 regulates miRNA biogenesis

We next examined mRNA levels of the known components in miRNA pathway in the mutant. Most of the components, including DCL1, HYL1 and SE itself as well as miRNA degrading enzymes (SDNs), displayed similar expression in *said1-3; said2-3* compared to Col-0 with the difference within 1.5-fold (**Figure S3C**). This result suggested that the increased miRNA accumulation in *said1-3; said2-3* was not caused by the de-regulation of the known genes in miRNA pathway, with a further suggestion that SAID1/2 might directly impact miRNA biogenesis. Of note, the transcript of AGO1 was accumulated in the mutant (**Figure S3C**), likely due to feedback regulation for increased need of RISC to house the accumulated miRNAs in the mutant.

Then, we assessed the steady-state level of pri-miRNAs. Whilst the recovered reads of pri-miRNA debris were low in RNA-seq, the overall perception was that numerous pri-miRNAs displayed reduced levels in *said1-3; said2-3* as observed in *chr2-1* relative to Col-0 (**Figure S4G**). The reduction of pri-miRNA levels could result from reduced transcription or more efficient processing of the transcripts in the mutants. Histochemical and western blot assays with GUS reporters for *MIR159* and *MIR164* loci showed that the transcription of the reporters might be marginally increased in the *said1-3; said2-3* background to their sibling controls (**Figures S4H and S4I**). These results implied that pri-miRNAs might be processed more efficiently in *said1-3; siad2-*3 vs Col-0. However, the results also indicated that SAID1/2 might inhibit the expression of certain *MIR* loci, and such regulation might contribute in part to reduced miRNA accumulation in the wild-type condition. How exactly SAID1/2 regulate transcriptional regulation will be studied elsewhere and not be discussed further here.

### SAID1/2 promote degradation and phosphorylation of SE

SAID1 and SAID2 did not seem to alter transcript levels of the known factors that regulate miRNA biogenesis and turnover. However, western blot assays indicated that protein accumulation of SE, AGO1 and possibly HYL1 was all increased in *said1-3; said2-3* compared to that of WT and the single *said1* and *said2* mutants (**Figure 4A**). Since SAID1/2 are IDPs, we hypothesized that they might bind SE and regulate its stability. To test this, we performed a cell-free protein decay assay using lysates extracted from three-week-old seedlings. When cycloheximide (CHX) was applied to block protein translation, SE had a half-life of approximately 20 min and was barely detectable afterwards in WT lysates. By contrast, SE protein was relatively more stable and clearly detected even in 60 min in the extracts from *said1-3; said2-3* (**Figure 4B**). In either case, the proteasome inhibitor MG-132 could inhibit SE decay, and SE protein levels were relatively stable and detectable in the extended incubation periods (**Figure 4C**). The results suggest that increased stability might account for the elevated SE accumulation in *said1-3; said2-3*, with further suggestion that SAID1/2 are negative regulators of SE accumulation.

**Figure 4.**
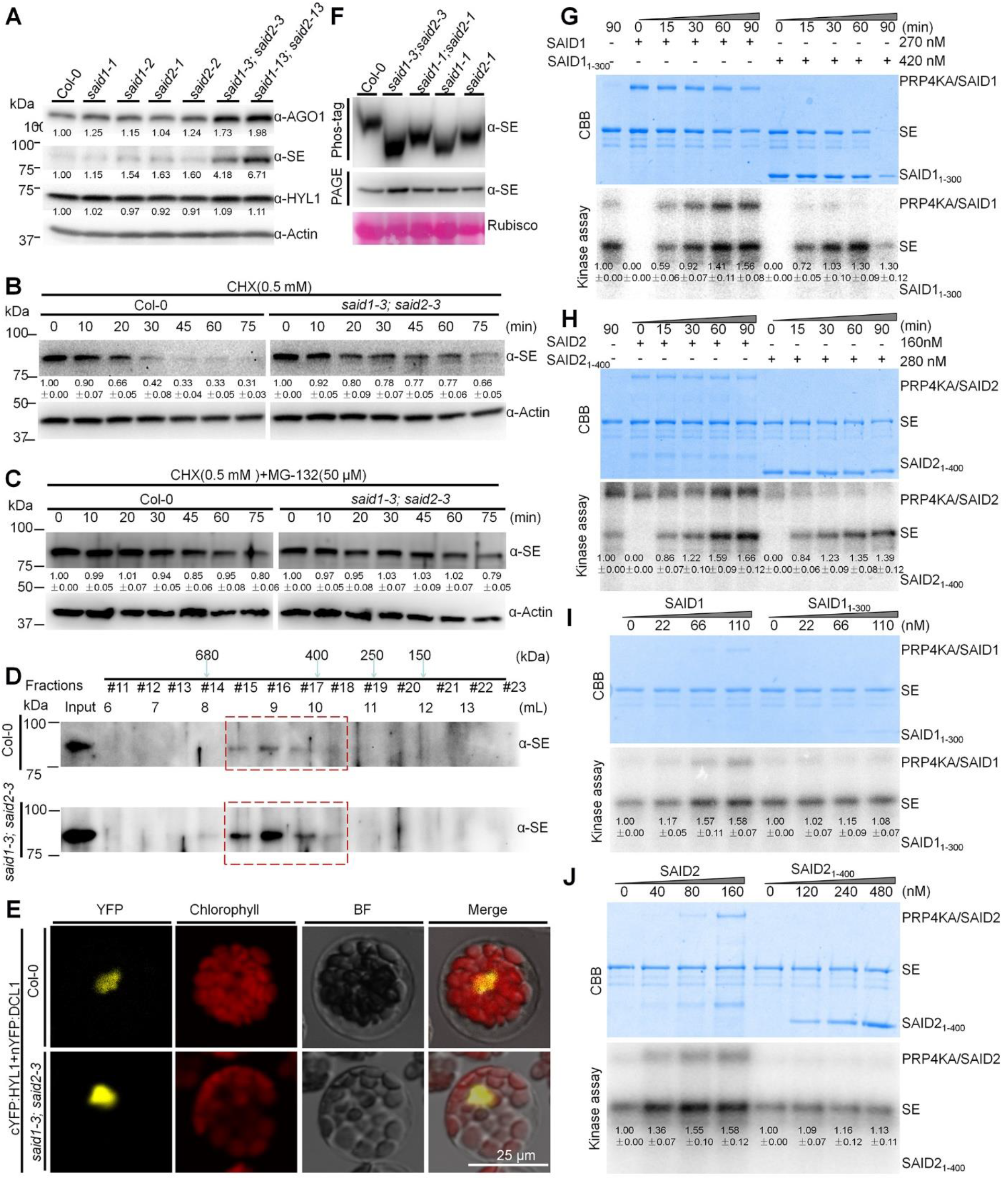
SAID1/2 promotes SE phosphorylation and degradation. (**A**) Western blot analysis showed the accumulation of AGO1 and SE in single and double mutants of *said1* and *said2* vs Col-0. Antibodies specifically recognizing AGO1, SE, HYL1 and Actin (loading control) were used. (**B-C**) Western blot analysis revealed that SE had extended half-life in cell-free extract from *said1-3; said2-3* than from Col-0. Total cell lysis prepared from the indicated lines was treated with CHX with or without MG-132 at the indicated time points before Western blot assays using an anti-SE antibody. Actin was a loading control. The numbers below the gels indicate the relative mean signals of SE at different time points that were normalized to those of SE at time 0, where the value was arbitrarily assigned a value of 1. (**D**) SEC revealed similar distribution spectra of SE protein in Col-0 and *said1-3; said2-3* plants. Total protein was extracted from 3-week-old *said1-3; said2-3* and Col-0 and resolved by the Superdex 200 10/300 GL column before western blot analysis using a specific anti-SE antibody. Elute fractions, volumes and corresponding molecular weight of protein standards were indicated. Dashed red boxes indicated the fractions with major elution of SE. (**E**) BiFC assays showed that the assembly of HYL1/DCL1-contained microprocessors was not inhibited in protoplasts prepared from *said1-3; said2-3* compared with Col-0. At least 20 protoplasts with similar signal were detected. Scale bars=25 µm. (**F**) Western blot analysis revealed that SE is hypo-phosphorylated in *said1; said2* compared to Col-0. SE was detected by SDS-PAGE and phosphor-tag gel in parallel using an anti-SE antibody. Rubisco served as loading control. (**G-J**) Kinase assays show that SAID1/2 promote SE phosphorylation by PRP4KA. Time-course (**G-H**) and dose-dependent (**I, J**) kinase assays of SAID1 (**G, I**) or SAID2 (**H, J**) impact on SE phosphorylation. Coomassie blue staining (**top panel**) and autoradiography (**bottom panel**) of indicated proteins were detected and quantified. The dosage of recombinant SAID1/2 and variants proteins were shown on the top of gels. Relative kinase activity with s.d. from three replicates is quantified. Represented results from three biological repeats are shown for panels (A to J)

We recently found that PRP4KA phosphorylates SE and triggers its degradation (Wang et al., 2022). We hypothesized that the phosphorylation status of SE might be altered in *said1-3; said2-3*. Indeed, phosphor-tag gel blot clearly showed the faster migration of SE protein from *said1-1; said2-1* and *said1-3; said2-3* relative to that of WT (**Figures 4F and S5A**). SE protein from the single *said1-1* and *said2-1* mutants also appeared to migrate faster but to less extent compared to that of *said1-3; said2-3* (**Figure 4F**). These results indicated that SE is hypo-phosphorylated in the mutants vs Col-0, with a suggestion that SE is stabilized in a hypo-phosphorylation form in the double mutants. We next surmised that SAID1/2 might directly interact with PRP4KA and/or PAG1 to regulate phosphorylation and degradation of SE. However, these possibilities were disapproved by Y2H assays (**Figures S6A and S6B**). We also performed Y3H assays to assess whether SAID1/2 promote the interaction of PRP4KA with SE, which might in turn facilitate the phosphorylation. Unexpectedly, addition of either SAID1 or SAID2 to the combination of PRP4KA and SE resulted in slower growth of yeast colonies vs the combination of PRP4KA/SE alone. This result superficially suggested that SAID1/2 might block PRP4KA/SE interaction. However, further western blot assays showed that SE accumulation was largely reduced once SAID1/2 were expressed (**Figure S6D**). This result indicated that SAID1/2 could trigger SE degradation in yeast similarly to in plants.

SAID1/2 stimulation of SE degradation in both Arabidopsis and yeast promoted us to hypothesize that the two IDPs might alter SE conformation in vivo, facilitating PRP4KA enzymatic activity on SE. To test this, we purified recombinant full-length SAID1/2 and their truncated variants SAID1_1-300aa_ and SAID2_1-400aa_ that lost interaction with SE. An in vitro kinase assay showed that addition of high-dosage of wild-type SAID1 (~270 nM) marginally impacted SE phosphorylation signals relative to SAID1_1-300aa_ **(Figures 4G and S5B)**. To our surprise, PRP4KA could also phosphorylate SAID1 but not its truncated mutant SAID1_1-300aa_ in vitro. This result suggested that two substrates might compete for limited amount of the kinase and ATP in the reaction system. To minimize the potential competition, we added a low dosage (up to 110 nM) of SAID1 or SAID1_1-300aa_ to the kinase assay system. Remarkably, SAID1 clearly boosted the phosphorylation signal of SE compared to SAID1_1-300aa_ **(Figures 4I and S5D)**. This result suggested that once mingled together, SE and SAID1 both become the substrates of PRP4KA and mutually promote the phosphorylation of each other in vitro. PRP4KA appeared to phosphorylate SAID2 in vitro as well. In this scenario, SAID2 clearly and progressively boosted SE phosphorylation signal whereas the SAID2_1-400aa_ did not (**Figures 4H, 4J, S5C, and S5E**). Altogether, both in vivo and in vitro assays showed that SAID1/2 could enhance PRP4KA activity on SE and promote its degradation in vivo.

### Accumulated SE does not affect the assembly of its scaffolded macro complexes

Homeostasis of SE accumulation is critical for its proper functions in different macromolecular complexes as excessive SE often causes assemblies of intermediate complexes and subsequently protein destruction as observed in *prp4ka* and *pag1* mutants (Li et al., 2020c; Wang et al., 2022). To test if this was the case in the *said1-3; said2-3* mutant, we performed size exclusion chromatography (SEC) assays. We observed that the distribution patterns of SE fractions were comparable between Col-0 and *said1-3; said2-3* with most of the SE fractions eluted at the 8-10 mL volume, equivalent to the range of 680-250 KDa (**Figure 4D**). This scenario was quite different from the broader ranges of SE elution in either *pag1* or *prp4ka* mutants (Li et al., 2020c; Wang et al., 2022). The contrasted results indicated that the accumulated SE protein in *said1-3; said2-3* does not seem to interfere with its assembly in macro-ribonucleoprotein complexes. To further test it, we co-expressed cYFP-HYL1 and nYFP-DCL1 in the protoplasts prepared from Col-0 and *said1-3; said2-3*. The YFP signal, indicative of interaction of the HYL1 and DCL1, was even stronger in *said1-3; said2-3* vs Col-0 (**Figure 4E**). The results validated that increased accumulation of SE did not disrupt the assembly of microprocessor complexes in the *said1-3; said2-3* mutant, likely because the accumulation of several tested factors in miRNA biogenesis and RISC effector were concertedly increased in the mutants (**Figure 4A**).

### SAID1/2 bind to pri-miRNAs

We further investigated if SAID1 and SAID2 impacted pri-miRNA processing. Electrophoretic Mobility Shift Assay (EMSA) showed that purified SAID1/2 did not bind to homogenous single-stranded (ssRNA) that comprised of 44nt poly(A)s (**Figures 5A and 5B**). To our surprise, SAID1/2 both could bind to hairpin structured pri-miRNAs with dissociation constants (Kds) of 1.950 ± 0.317nM, and =0.996 ± 0.179nM, respectively (**Figure 5C-5E and S7A-C**). Since the truncated variants of SAID1/2 barely associated with pri-miRNAs, or bound pri-miRNAs with much reduced affinity (**Figures 5F, S7D, and S7E**), then potential RNA binding domains are likely located in the middle and C-terminal parts of the proteins. Importantly, we conducted in vivo RIP assays and observed that SAID1/2 immunoprecipitates (IP) were clearly enriched with selected pri-miRNAs relative to a control IP (**Figure 5G**). Thus, both in vitro and in vivo assays showed that SAID1/2 are dsRNA-binding proteins.

**Figure 5.**
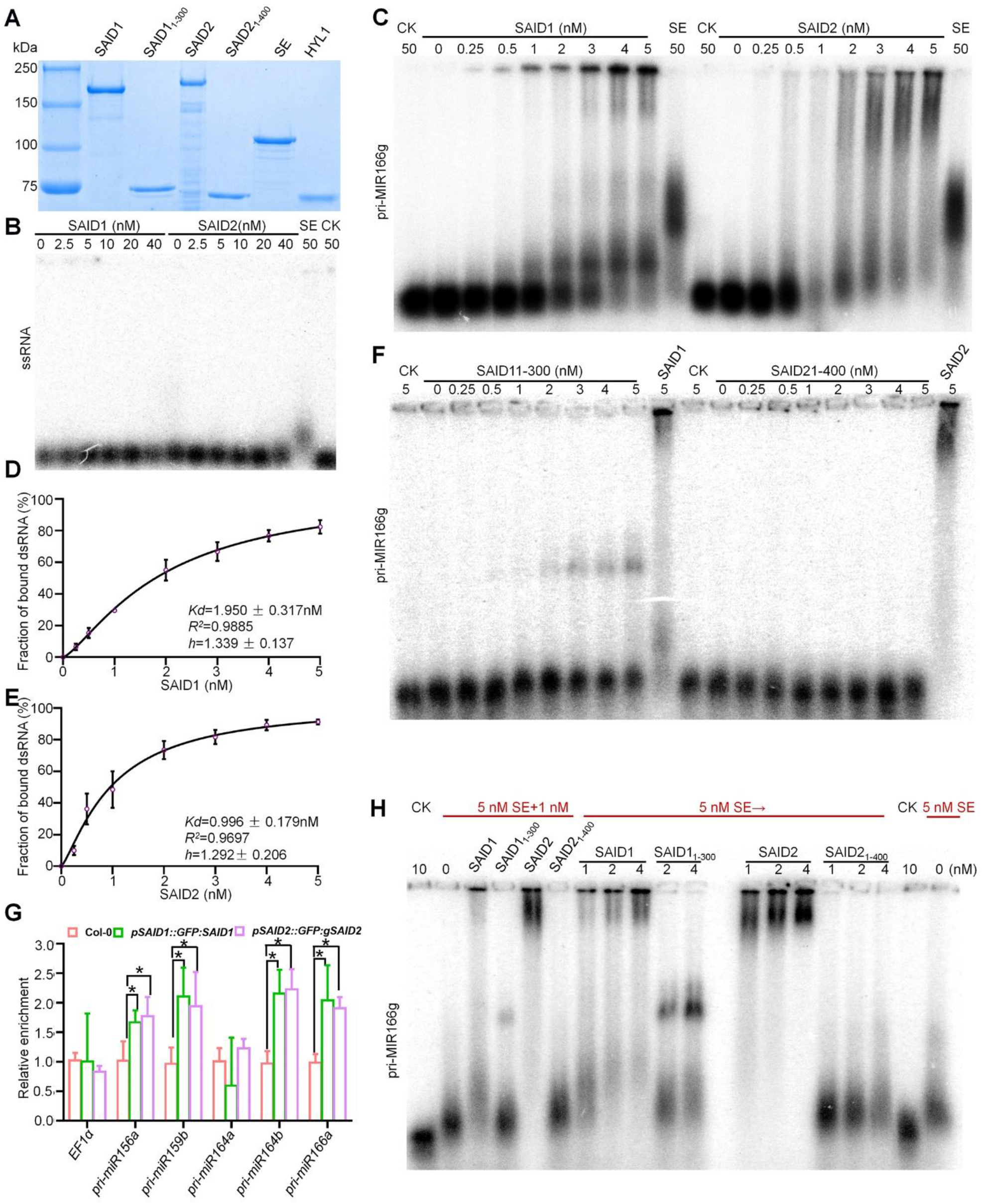
EMSA shows that SAID1/2 have high binding affinity to pri-miRNAs. (**A**) Coomassie blue staining of recombinant SE, HYL1, full-length and truncated SAID1/2 proteins from *E. coli* or a baculovirus-insect system in SDS-PAGE. (**B, C**) EMSA shows the mobility patterns of SAID1/2 with 44 nt Poly(A) ssRNA (**B**) and pri-miRNA (**C**). His-SUMO (CK), and 50 nM of SE served as negative and positive controls, respectively. Protein concentrations are shown above. (**D, E**) The binding affinities (Kds) of SAID1 and SAID2 with pri-miRNA, calculated based on EMSA mobility of (**C**). Data are means ± SE of three biological replicates. (**F**) EMSA shows the poor binding affinity of truncated SAID1/2 variants with pri-miRNA. (**G**) RIP assays show that SAID1/2 bind to pri-miRNAs in vivo. The relative signal of pri-miRNAs was calculated with s.d. from three biological repeats (*P < 0.05; **P < 0.01; unpaired, two-tailed Student’s *t*-test). (**H**) Competing binding assays of SE and SAID1/2 with pri-miRNA. Mobility pattern of RNA-protein after co-incubation of SE and SAID1/2 with pri-miRNA166g (lanes 3-6, indicated as SE+), or sequentially incubation of SE with pri-miRNA166g followed by incubation with various concentrations of SAID1/2 (indicated as SE→). His-SUMO served as negative control (CK).

The Kd(s) of SAID1 and SAID2 association with pir-miRNAs are at a similar scale of that of HYL1 (Kd= 0.7 nM) but much less than that of SE (Wang et al., 2018), indicating that SAID1/2 have binding affinity with pri-miRNAs comparable to that of HYL1, but retain significantly higher affinity with pri-miRNAs than SE, with a further suggestion that SAID1/2 might sequester pri-miRNAs from SE protein. To test this, we pre-incubated SE with pri-miRNA followed by addition of various concentrations of SAID1/2, or their truncated variants (**Figure 5H**). EMSA results showed that addition of the truncated variants did not alter the migration pattern of SE/pri-miRNA complex, whereas the electrophoretic mobility of pri-miRNA-protein complexes either recapitulated that of pri-miRNA-SAID1/2 or became slower than that of SE-pri-miRNAs (**Figure 5H**). These results indicated that the addition of full-length SAID1/2 proteins indeed sequestered pri-miRNAs from SE protein. Similar results were also obtained when the order of protein incubation with RNA was reversed (**Figure S7F**). Taken together, the results further validated that SAID1/2 retained higher binding affinity with pri-miRNAs than SE and could readily sequester/occupy pri-miRNAs from SE to prevent the downstream processing.

### SAID1/2 inhibit pri-miRNA processing

Given the fact that SAID1/2 can sequester pri-miRNAs from SE and the fact that *MIR* loci are often de-repressed while the steady-state levels of pri-miRNAs are reduced in *said1-3; said2-3*, miRNA production might be more efficient in the mutant vs WT. To address the direct impact of SAID1/2 on pri-miRNA processing, we performed *in vitro* microprocessor reconstitution assays (Wang et al., 2018). First, we pre-incubated either SAID1 and SAID2, or their truncated variants, with pri-miR166f and pri-miR166g for 15 min before addition of SE and the core microprocessor component DCL1/HYL1 (**Figures 6A and 6D**). We observed that supply of truncated SAID1/2 exerted minor or literally no effect on pri-miRNA processing (**Figures 6B, 6C, 6E, and 6F**). By contrast, pre-incubation of either SAID1 or SAID2 with pri-miRNAs greatly decreased the pri-miRNA processing efficiency, and this inhibition was exerted in a dose-dependent manner *in vitro* (**Figures 6 B, 6C, 6E, 6F, and S8A-D**). We also pre-incubated SE with pri-miRNA before addition of SAID1/2 or their truncated variants as the prevailing model is that SE binds to pri-miRNAs co-transcriptionally (Gonzalo et al., 2022; Zhu et al., 2022). Again, SAID1/2, but not the SE-interaction compromised mutants, still inhibited microprocessor activity. Even more, the inhibition might be even more severe compared to the situation when SAID1 and SAID 2 is pre-incubated with pri-miRNA earlier than SE (**Figures S8E and S8F**). One possibility is that pre-incubation of SE with pri-miRNAs might alter the conformation of the ribonucleoprotein complex and facilitate its recruitment of SAID1/SAID2 for inhibition. This scenario might imply that SE recruits SAID1/2 that in turn sequester pri-miRNAs from SE to inhibit pri-miRNA processing in vivo.

**Figure 6.**
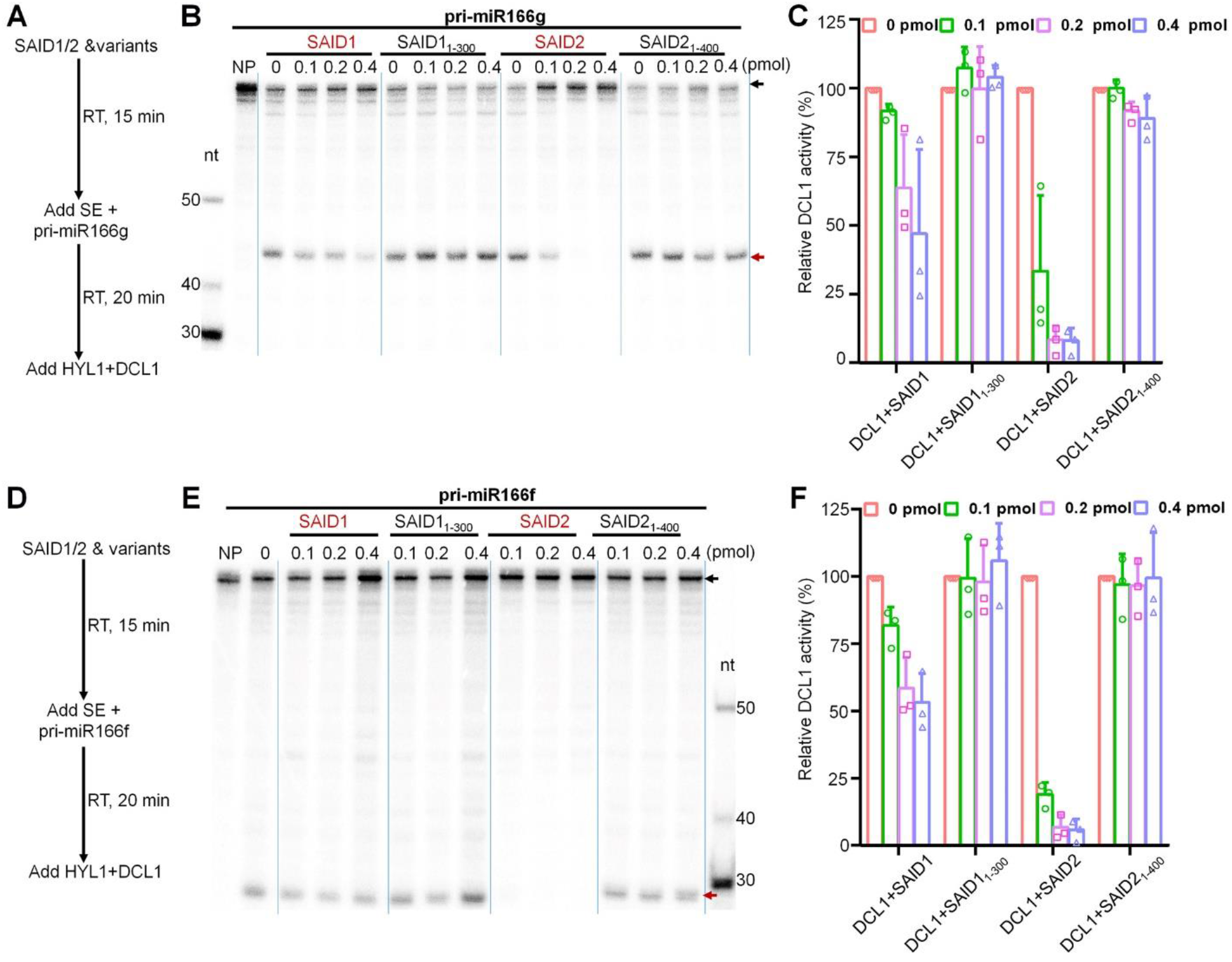
SAID1/2 inhibit pri-miRNA processing in microprocessor reconstitution assays. (**A, D**) Schematic illustration of reconstitution orders of microprocessor components *in vitro*. (**B, C, E and F**) SAID1/2 significantly inhibited the processing efficiency of pri-miR166g (**B, C**) and pri-miR166f (**E, F**) *in vitro*. In (**B, E**) autoradiography of in vitro microprocessor assays. Black and tangerine arrows indicate processed and unprocessed fragments of pri-miRNAs. Various concentrations of SAID1/2 or their truncated variants were subsequentially incubated with the reconstituted microprocessor and 1,000 cpm (counts per minute) of 5’ labeled pri-miR166g and pri-miR166f with [γ-^32^P] ATP. (**C, F**) Quantification of relative cleavage efficiency with s.d. from three replicates.

### SAID1/2 dysfunction do not alter LLPS capacity of SE and HYL1

To further investigate how SAID1/2 regulate SE function in vivo, we performed a serial of confocal experiments (**Figures 7 and S9**). SE is an IDP and has been reported to show a feature of LLPS when expressed under a constitutive promoter (Xie et al., 2021). In our hands, SE-tagged mCherry could indeed form numerous condensates or foci in nucleus when transiently expressed under *Super* promoter (**Figure 7E**). However, SE-GFP expressed in the native promoter in the transgenic lines appeared to be distributed homogenously in nucleus. A few tiny nuclear sparkles might be observed from time to time, but the sizes were typically smaller than the ones when overexpressed (**Figures 7A, 7C, and 7E**). When the small particles were beached by laser, the GFP fluorescence could be recovered within 20s, suggestive of LLPS feature (**Figure 7C**). Since the majority of SE-GFP is evenly dispersed in nucleoplasm (**Figure 7A and 7C**), we used HYL1-GFP as a proxy of D-bodies. Nucleus-localized HYL1-GFP displayed 1-2 bright foci in the root tip of stable transgenic plants. While most cells in WT showed 1 HYL1-GFP foci in their nuclei, increased number of cells in *said1-3; said2-3* showed 1 or 2 brighter HYL1-GFP foci in the nuclei, indicative of efficient D body assembly in the mutant (**Figure 7B**).

**Figure 7.**
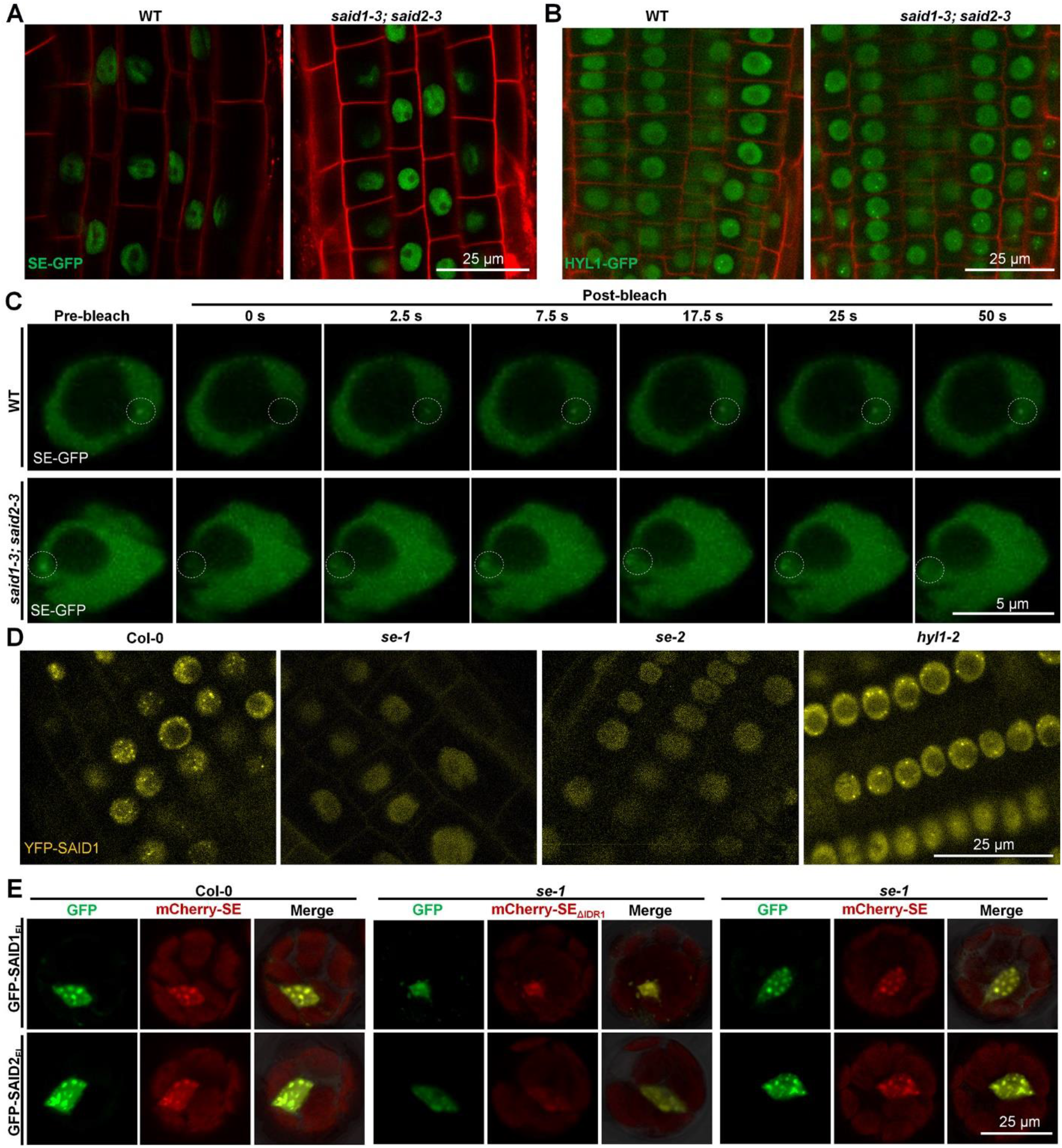
SAID1/2 nucleate on SE to form condensates. (**A, B**) Expression pattern of SE-eGFP (**A**) and HYL1-eGFP (**B**) in root tip of seven-day-old transgenic plants expressing *pSE::gSE:eGFP* or *pHYL1::gHYL1:eGFP* in Col-0 and *said1-3; said2-3* background. Scale bars=25µm. Note: elevated eGFP-SE signal (**A**) and increased numbers of cells with 1 or 2 HYL1-eGFP speckles (**B**) were observed in *said1-3; said2-3* vs col-0. (**C**) FRAP assay of SE-eGFP in root tips of 7-day-old homozygous WT and *said1-3; said2-3*. FRAP of SE-eGFP was analyzed in RAM of 7-day-old homozygous *pSE::gSE:eGFP* transgenic plants in Col-0 and *said1-3;said2-3* background. White circles indicate the regions of interest (nuclei speckles) that were bleached. Scale bars=5 µm. Images are representative of at least 10 nuclei for each genotype in three independent assays. Note: GFP signal was quickly recovered after the bleaching was stopped. (**D**) The eYFP-SAID1 signal detected in root apical meristems in 7-day-old seedlings of homozygous *pSAID1::eYFP:gSAID1* in *se-2* or *hyl1-2* backgrounds. Scale bars=25 µm. (**E**) Nuclear condensation of eGFP-SAID1/2 in mesophyll protoplasts of Col-0 and *se-1* mutants. *Super:eGFP-SAID1/2* were co-transformed with either *Super:mCherry-SE* or truncated *Super:mCherry-SEΔ*_*IDR1*_ into the protoplasts. Scale bars=25 µm. Representative images of at least 10 nuclei for each genotype in three independent assays are shown.

We also assessed effects of SAID1/2 mutation on the assembling of D bodies and LLPS of SE-GFP, no significant differences were observed between WT and *said1-3; said2-3* mutants (**Figure 7C**). Phase separation capacity of HYL1-GFP was also not affected in *said1-3; said2-3* mutants (**Figure S9**), indicating that microprocessor assembling and LLPS of SE was not interfered and was maintained normal in *SAID1*/*SAID2* mutants vs Col-0.

### SAID1/2 display LLPS feature and regulate SE function in the condensates in vivo

SAID1/2 are also predicted to be IDPs with several intrinsic disordered regions identified (**Figures S10A-F**). We also examined whether SAID1/2 could form nuclear condensates. When expressed transiently in protoplast, full length protein of SAID1/2 were detected in the nucleus, and clearly assembled in the nuclear condensates (**Figures 7E, S10G, and S10H**), reminiscent of SE foci in the overexpression lines (Xie et al., 2021). We generated stable transgenic plants expressing native promoters-driven YFP-SAID1 and GFP-SAID2. Both SAID1 and SAID2 expressed ubiquitously at the root, hypocotyl, cotyledon, true leaves and shoot apical meristem among other tissues (**Figure S11A and S11C**), indicating their ubiquitous functions in regulating plant development and adaption. SAID1-YFP, SAID1-GFP and SAID2-GFP were localized in the nucleus, and they also assembled into nuclear condensates (**Figures 7D, 7E, and S11**).

Importantly, they showed phase separation patterns, as evidenced by fast recovery in FRAP assays of GFP-SAID1 and GFP-SAID2 (**Figure S10I**). Next, we tested whether the IDRs were crucial for their capacity to form nuclear condensates by examining expression pattern of truncated SAID1/2 with depleted IDRs. Compared with full-length SAID1, fluorescent signals of the truncated SAID1 proteins (SAID1_Δ5-55aa_, SAID1_Δ377-407aa_, SAID1_Δ377-580aa_, and SAID1_Δ515-577aa_) was decreased, additionally, the number and size of nuclear condensates were also decreased, but not completely disrupted (**Figure S10G**). As for SAID2, depletion of amino acid 656-727 of SAID2, but not amino acid 33-77 resulted in reduced protein level, less number and smaller size of nuclear condensates (**Figure S10H**). The results showed that SAID1/2 are nuclear-localized protein and assembled in the nuclear condensates, and their stability and assembling in nuclear condensates rely on their IDRs.

Next, we investigated SAID1/2 expression pattern in the *se* mutant background. When expressed in protoplast, dimmer signal of SAID1/2 was detected in *se-1* vs Col-0. Meanwhile, no obvious nuclear condensates of SAID1/2 were detected in the *se-1* protoplast compared with that of Col-0. Introduction of mCherry-SE_ΔIDR1_ into *se-1* did not restore the nuclear condensation of SAID1/2. By contrast, mCherry-SE could recover the nuclear condensation of GFP-tagged SAID1/2 (**Figure 7E**). Finally, we examined expression patterns of native SAID1/2 in Col-0 and *se* mutants. Again, much faint signal of eYFP-SAID1 or eGFP-SAID2 was detected in *se-1* and *se-2* mutants, whereas strong signal and nuclear condensates of eYFP-SAID1 or eGFP-SAID2 were detected in Col-0 background (**Figures 7D and S11E**). By contrast, the nuclear condensation patterns of SAID1/2 were minimally altered in *hyl1* vs Col-0 (**Figures 7D and S11E**). These results indicated that SAID1/2 nucleate on SE to form condensates, which in turn promote SE degradation or deprive pri-miRNAs from SE to inhibit pri-miRNA processing in vivo.

## Discussion

SAID1/2 are annotated as homologs of DSPP in metazoans and theoretically involved in depositing mineral materials to extracellular matrixes to harden cells. Unexpectedly, we here identified SAID1/2 as novel partners of SE and their inhibitory roles in miRNA biogenesis. We proposed a model that SAID1/2 act as dual inhibitors of SE function in miRNA biogenesis.

Several pieces of evidences supported our notion: First, SAID1/2 physically interact with SE (**Figures 1A-F and S1**), and their interaction has been clearly validated by multiple assays and by independent laboratories (Bajczyk et al., 2020); Second, SAID1/2 promotes SE phosphorylation in vivo and in vitro, triggering its destruction in plants (**Figures 4 and S5**); Third, SAID1/2 possess very strong binding affinity to pri-miRNAs and can sequester the substrates from SE protein which only has moderate binding affinity to pri-miRNAs (**Figures 5 and S7**). Finally, SAID1/2 could directly inhibit pri-miRNA processing by microprocessor (**Figures 6 and S8**). Given that the SAID1/2 condensate on SE in nucleus (**Figures 7D, 7E, and S11E**), we envision that the two proteins could hijack pri-miRNAs from SE to prevent the downstream microprocessor activity while the proteins could also stimulate phosphorylation of SE via PPR4KA, leading to degradation of SE, so that miRNA production is further stringently confined in plants (**Fig. 8**). Thus, SAID1/2, represents a unique mode of action of controlling SE function that takes places at the levels of posttranscriptional processing and posttranslational modification. This mode of action is different from that of CHR2 that can sequester pri-miRNA and remodel the RNA secondary structure to inhibit miRNA production (Wang et al., 2018).

**Figure 8.**
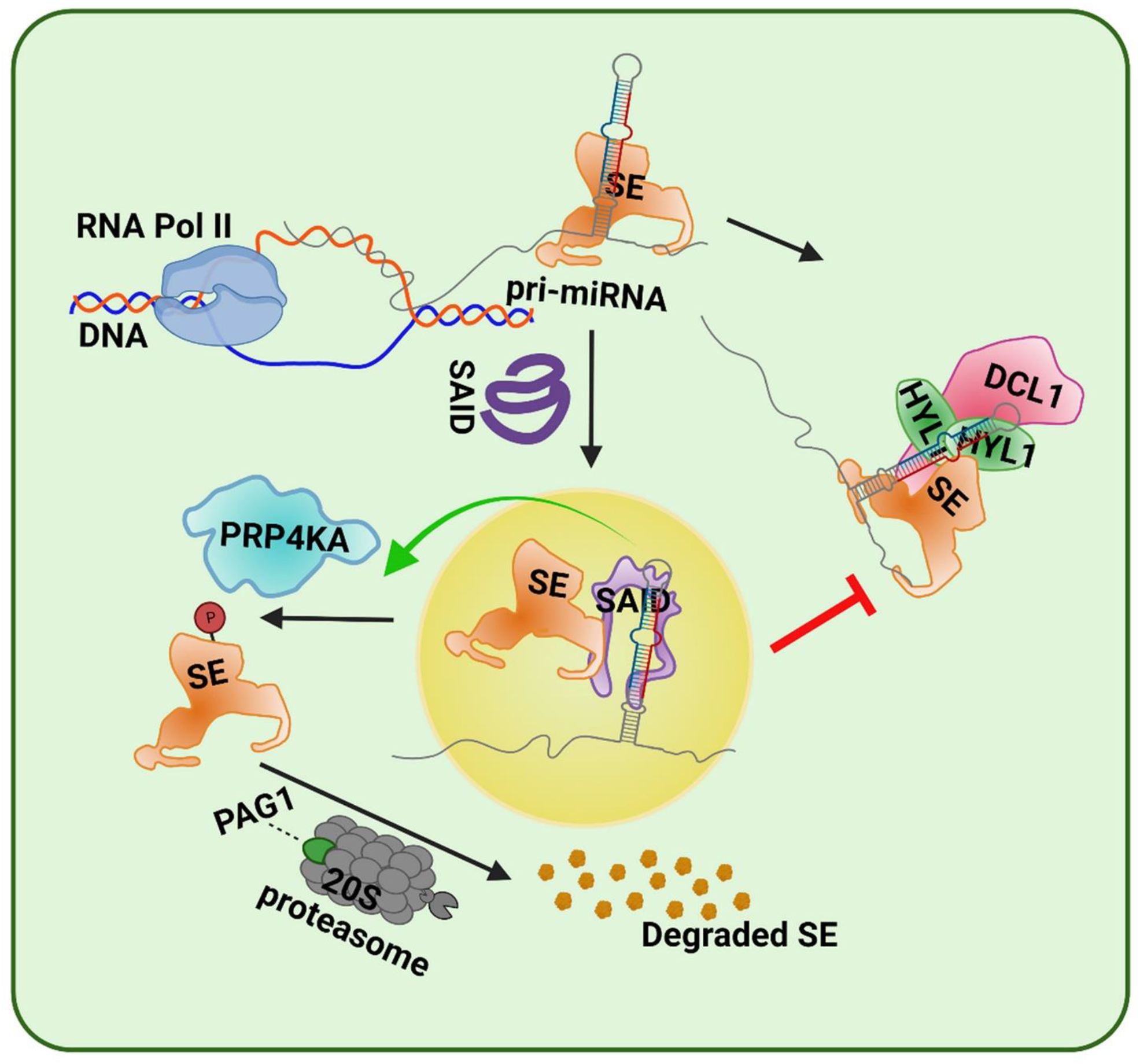
Proposed model for dual inhibitory roles of SAID1/2 on SE functions in miRNA production. The IDP SAID1/2, which possess LLPS characteristics, form nuclear condensates on SE (highlighted in yellow cycle), deprive pri-miRNAs from SE to prevent the downstream microprocessor activity while promoting SE phosphorylation and resultant destruction. Thus, SAID1/2 dually inhibit SE function to fine-tune miRNA expression. Once SE is reduced, SAID1/2 condensation is disassembled to release the inhibition of SE. Thus, SAID1/2 and SE form a feedback regulatory circuit.

SAID1/2 are also distinct from a recently reported IDP, Constitutive Alternations in the Small RNAs pathway 9 (CARP9) that promotes RNA silencing by facilitating shuttling of mature miRNA from HYL1 to AGO1-mediated RISC (Tomassi et al., 2020).

### Mechanisms for dual inhibitions of SAID1/2 on miRNA production

How could the same proteins exert dual inhibitory functions on SE? The dual inhibitions might result from the unique biochemical features of SAID1/2; and might also be independent from each other. SAID1/2 proteins are IDPs, and so is SE. Once attached to SE, SAID1 or SAID2 might change the conformation or configuration of SE, exposing its serine or threonine residues to the kinase PRP4KA, and facilitating the catalytic activity. Alternatively, SAID1/2 might directly bind and channel adenosine triphosphate (ATP) and facilitate the transfer of a phosphate moiety from ATP to the substrate molecules. Additionally, SAID1 and -2 might also bind and store ADP to enforce the catalytic reaction forward. The last scenario might explain the observation that SAID1/2 could also stimulate phosphorylation of SAID1/2 by PRPK4A in vitro, although they seem not to interact with each other in vitro (**Figures 4G-J, S5, and S6**). For in vivo, SAID1/2-mediated promotion of SE phosphorylation likely involves LLPS as the SAID1/2-SE nuclear condensates might serve as a platform to recruit PRP4KA for enzymatic activity (**Figures 7, S10, and S11**).

SAID1/2 inhibitory effect on microprocessor activity is likely through its high affinity of pri-miRNAs. The two proteins have predicted isoelectric points as 6.94 and 7.64, respectively, and literally neutral in a physiological condition. Thus, the possibility of the non-specific package of pri-miRNAs by the two proteins can be excluded. Further supporting of this view are two facts that Hill parameters of SAID1/2 association with pri-miRNAs are close to 1 (the molarity ratio of protein: RNA) and that SAID1/2 do not bind to ssRNA (**Figures 5B-5H and S7**). Thus, the association of SAID1/2 with pri-miRNAs likely involves the recognition of RNA secondary structure. Of note, the two proteins do not appear to have classic RNA binding or recognizing motif (RBM or RRM). Deep-learning predicts that SAID1/2 possess spaghetti-like 3D structures centered on small alpha-helix domains. It is possible that upon supply with pri-miRNAs, the IDPs might re-shape their structures, which in turn enable the accommodation of pri-miRNAs with high affinity. This scenario is reminiscent of the discovery of FVE, the protein which harbors WD domains and would otherwise be thought to be only involved in protein-protein interaction, does not have canonical RRM or RBM but can also bind to double-stranded RNA with strong affinity (Sun et al., 2021).

It did not escape from our attention that SAID1_1-300_ retains certain binding capacity of pri-miRNAs but does not interact with SE (**Figures 5F, S7D, and S7E**). Whereas SAID1 represses pri-miRNA processing significantly, truncated SAID1_1-300_ does not (**Figures 6 and S8**). The results imply that SAID1 inhibition of pri-miRNA processing entails its physical interaction with SE. Taken together, the novel IDPs SAID1/2 are recruited to SE scaffolded ribonucleoprotein complexes, stripping pri-miRNAs from SE, and inhibiting pri-miRNA processing, while the proteins could also promote PRP4KA-mediated phosphorylation of SE to elicit its degradation (**Fig. 8**).

### Physiological significance for SAID1/2 repression of SE function

miRNAs are crucial regulators of gene expression in eukaryotes and their accumulation is spatiotemporal changes in response to interior physiological stimuli and exterior environment. SAID1/2-mediated double inhibition of SE function might provide a gateway to quickly change miRNA accumulation, which could further re-program transcriptome for better adaption to physiological changes. Indeed, the accumulation of numerous miRNAs is restricted into certain niches, whereas the expression of many others needs to be reprogrammed in response to environmental stresses and stimuli (Li et al., 2017; Song et al., 2019). Thus, proper adjustment of protein accumulation of microprocessor components including SE, a master regulatory cohort for RNA metabolism might be an efficient checkpoint. Under these scenarios, accumulated SAID1/2 could promote SE phosphorylation via PRP4KA and facilitate its degradation, leading to reduced miRNA levels. Conversely, lowered expression of SAID1/2 would promote SE accumulation, and resultant miRNA upregulation. Furthermore, the accumulation of pri-miRNAs and miRNAs are often asynchronized. In numerous cases, expression of pri-miRNAs increases but miRNAs are downregulated, and vice versa (Barciszewska-Pacak et al., 2015; Choi et al., 2020). As a strong RNA binding protein, SAID1/2 could serve as an insulator to separate pri-miRNA accumulation from miRNA production. For instance, SAID1/2 could simply sequester excessive pri-miRNAs from SE, preventing the downstream processing, thus miRNA homeostasis is maintained. Otherwise, lack of SAID1/2 would allow prompt handover of pri-miRNAs from SE to DCL1/HYL1 for processing. Indeed, the expression of SAID1/2 displays ubiquitous expression patterns but shows various fluctuations response to plant hormone treatments and light stimuli and circadian oscillation. Their expression is also de-regulated under different biotic (i.e., *Pseudomonas syringae*) and abiotic stresses such as cold, salt, osmatic, genotoxic and UV-B, and repressed by heat stress. The dynamic response of SAID1/2 expression to both developmental and stresses inferring their surveillance roles on expression homeostasis of SE and supply chains of pri-miRNAs. Thus, the dual inhibitory roles of SAID1/2 on SE serves as a prompt regulatory mechanism to fine-tune miRNA accumulation in responses to physiological and environmental variations.

### Inter-dependent and independent functions of SAID1/2 and SE

Whereas SAID1/2 have clear crosstalk with SE in miRNA pathway, they might each have their own regulatory regimes. This surmise can be highlighted by the fact that a majority of *said1/2*-deregulated genes are not overlapped with those in *se-2*, despite more than half of *se-2*-regulated genes fall into the territory controlled by *said1-3; said2-3* (**Figures 2A-C**). These scenarios might be accredited to biochemical features of the IDP proteins. Typically, IDPs partner with different scaffold proteins and perform various functions accordingly. It is highly likely that SAID1/2 might like SE contribute to assembly of numerous ribonucleoprotein complexes, participating in different regulatory processes of gene expression. In lines with this prediction, SAID1/2 could regulate transcription of a few tested *MIR* loci (**Figures S4H and S4I**). Also, SAID2 shows auto-activation in Y2H assays (data not shown), Thus, the proteins might be transcription factors, and particulate in regulation of genome-wide transcription. Whether SAID1/2 and SE coordinate to transcriptional regulation await future investigation. Additionally, SAID1 and SAID2 are remote paralogs in phylogenic tree, indicating that they might have functional specification.

Intriguingly, the genetic pathways responding to multiple hormones (i.e., salicylic acid) and auto-immunity have been upregulated (**Figures 2D-2G**), whereas general biological processes (such as photosynthesis, metabolite accumulation, cell wall biogenesis) are repressed in *said1; said2*, indicating that SAID1/2 proteins might control the balances of plant growth and defense in normal conditions. Loss-of-function mutants of *said1; said2* display short-internodes, more branches, and early flowering, but without clear defect in overall seed yield. These ideal agronomic traits likely result from sophisticated and mutual regulations of miRNA with phytohormones(Li et al., 2020b), immunity (Wang et al., 2019), and environmental stimuli (Li et al., 2017). These features make SAID1/2 ideal targets for genetic engineering of crops for better agronomic improvement. To a broader level, whether DSPP in metazoans act as plant SAID1/2 to regulate RNA silencing or immune responses, in addition to their classic roles in dentin hardening, remains to be explored.

## ACKNOWLEDGMENTS

We thank members of the Zhang lab for proofreading of the manuscript and Dr. S.W. Yang for anti-SE, anti-HYL1 antibodies. We thank Drs. Stanislav Vitha and Malea Murphy for their training, and Texas A& M University Microscopy and Imaging Center Core Facility (RRID: SCR_022128), Texas A& M University Integrated Microscopy & Imaging Laboratory for providing the microscopy facilities (RRID:SCR_021637). This work was supported by grants from NIH (R01GM127742 and R01GM132401) to X. Z.

## Data availability

The data generated during this study will be deposited in GEO once the manuscript is accepted.

## Author Contribution

X.Z. conceived the project. B.S., and X.Z., designed the experiments. B.S. performed most of the experiments and analyzed majority of the data. L.W., performed protein decay assay, kinase assay, and drew the model, X.Y., performed protein purification and participated in the in vitro microprocessor assay. Y.L., performed sRNA-seq library construction, guided RNA-seq library preparation, western blot, and sRNA blot. Y.L., B.S., and T.Z., performed analysis of sRNA-seq data. C.L., and B.S., performed analysis of RNA-seq data. Y.L., T.W., Zh.W., and ZY.W., initially started the project, generated partial of genetic materials and provided intellectual advice. C.W., contributed to Y2H and performed Y3H assay. X-G., S.C., T.O., C.T., X-Z., and X.P., contributed to plasmid constructions, antibodies, among other reagents and offered guidance and suggestion through the study. B.S. wrote the initial draft of manuscript and X.Z thoroughly edited the paper, all authors contributed to the proof-reading of manuscripts.

## Materials and Methods

### Plant materials and growth conditions

The *said1-1* (SALK_080762C), *said1-2* (SALK_133343), *said2-1* (SALK_062434), *said1-2*(SAIL_88_D02), *se-1* (CS3257), *se-2* (SAIL_44_G12), and *hyl1-2* (SALK_064863) mutants were obtained from the ABRC and verified by PCR analyses. The *said 1-1; said 2-1* T-DNA insertion double mutant was generated in this study by crossing *said 1-1* with *said 2-1*, and F2 homozygotes were isolated by PCR analyses. The *CRISPR-Cas9* null alleles of *said 1* and/or *said 2* were generated and validated via sequencing the PCR products of *SAID1* and *SAID2*. In the representative line (*SAID1*; *SAID2-Cas9 #3*), an insertion of 1 bp at +36 nucleotide (nt) of the *SAID1* CDS was detected, resulting in a pre-mature stop codon at the downstream 20 amino acid (aa) residues. By contrast, a deletion of 1 bp at the +2283 of the *SAID2* CDS was detected, leading to a pre-mature stop codon at the 784 aa residues. Next, the *SAID1*; *SAID2-Cas9 #3* line was crossed with Col-0 to clean the background, and the F3 generation line free of Cas9 insertion via PCR genotyping was used for further analysis.

The *pMIR159a::FM-GUS, pMIR159b::FM-GUS, pMIR164a::FM-GUS*, and *pMIR164b::FM-GUS* marker lines have been described previously (Wang et al., 2018), and the relative F3 homozygous lines were generated by crossing with *said1-3; said2-3* and verified by sequencing and PCR analyses in this study. The *pSAID1::eYFP:gSAID1, pSAID2::eGFP:gSAID2* transgenic lines, and the corresponding F3 homozygous lines were generated by crossing with *se-1, se-2*, and *hyl1-2* were generated and verified by PCR analyses in this study. The *pHYL1::gHYL1:eGFP* and *pSE::gSE:eGFP* marker lines were generated according to previous report (Fang and Spector, 2007), and the corresponding F3 homozygous lines were generated by crossing with *said1-3; said2-3* and were verified by sequencing and PCR analyses in this study. Unless otherwise specified, the Arabidopsis ecotype Columbia-0 (Col-0) served as WT in this study, and mutants in other background were crossed with Col-0 three times to clear the background. Seeds were sterilized and germinated on Murashige and Skoog (MS) medium after stratification for 3-4 days at 4 °C. Seedlings were grown horizontally on MS medium and plants were grown in growth chamber under photoperiod of 12-h light/12-h dark at 22±1 °C.

### Plasmid construction and plant transformation

CDSs or genomic DNAs were cloned into pENTR/D-TOPO (Thermo Fisher) or modified pENTR1A vectors and confirmed by sequencing. The binary vectors were generated by Gateway cloning or NEBuilder HiFi DNA Assembly as previously described (Wang et al., 2018). The primers used are listed in Supplementary Table 2. For the construction of pENTRY clones, *SAID1* and *gSAID1* were ligated to pENTR/D vectors, *SAID2*, truncated variants of *SAID1* and *SAID2* were ligated to Xcm1-digested pENTR1A vectors by T4 DNA ligase.

Vectors for Y2H were generated by Gateway attL-attR (LR) recombination reaction (Thermo Fisher) of modified *pGBKT7-DC BD* and *pGADT7-DC AD* with the above entry vectors ligated with *SAID1, SAID2* and relative truncated variants. Entry vectors were linearized with MluI (for pENTR/D vectors) or ApaI (for pENTR1A vectors) to disrupt Kanamycin resistance of entry vectors before LR with *pGBKT7-DC BD* empty vectors. For Y3H vectors by one-step cloning, SE was cloned into the EcoRI/BamHI-digested *pBridge-BD-MCSII* vector to yield *pBridge-BD-SE-MCSII. SAID1* and *SAID2* were cloned into the NotI/AscI-digested *pBridge-BD-SE-MCSII* to yield *pBridge-BD-SE-SAID1/pBridge-BD-SE-SAID2* final vectors. Vectors for BiFC were generated by LR of *pBA-nYFP-DC* and *pBA-cYFP-DC* with the entry vectors ligated with *SAID1, SAID2*.

To construct *pBA002a-pSAID1::eYFP:gSAID1*, promoters of *SAID1* (3366bp) were cloned and ligated to EcoRV/XbaI-digested pBA002a-eYFP-DC vectors by one-step cloning, and *gSAID1*-pENTR/D was assembled with the modified vectors above by Gateway attL-attR (LR) recombination reaction (Thermo Fisher) to yield final binary vectors. To construct *pCAMBIA1300-pSAID1::eGFP:SAID1, SAID1* were cloned and ligated to BamHI/KpnI-digested *pCAMBIA1300-eGFP-MCS* vector by one-step cloning, and then the promoter of *SAID1* (3366 bp) was cloned and ligated to SalI-digested *pCAMBIA1300-MCS:eGFP-SAID1* by one-step cloning. To construct *pCAMBIA1300-pSAID2::eGFP:gSAID2, gSAID2* were cloned and ligated to BamHI/KpnI-digested *pCAMBIA1300-eGFP-MCS* vector, and the promoter of *SAID2* (2457 bp) was cloned and ligated to SalI-digested *pCAMBIA1300-eGFP-gSAID2* by one-step cloning. *pCAMBIA1300-pHYL1::gHYL1:eGFP* and *pCAMBIA1300-pSE::gSE:eGFP* were generated according to previous report (Fang and Spector, 2007), promoter and genomic DNA were cloned and ligated to SalI/XbaI-digested *pCAMBIA1300-MCS*-*eGFP* by one-step cloning.

To construct *SAID1/SAID2-CRISPR-Cas9*, guide sequences targeting *SAID1/SAID2* were designed. Guide sequences and corresponding complementary strands were annealed, BsaI-digested, and ligated with BsaI-digested *AtU6-26–sgRNA* vector by T4 DNA ligase. Tandem sgRNA sequences of *SAID1* sgRNA and *SAID2* sgRNA were obtained by ligating SpeI-digested *SAID2* sgRNA with SpeI/NheI-digested *AtU6-26-SAID1 sgRNA*. The tandem sgRNA sequences were generated from relevant SpeI/NheI-digested *AtU6-26-sgRNA* vectors, and then were ligated to SpeI-digested *pCAMBIA1300-pYAO-Cas9-MCS* vector.

To generate *pAcGHLT-GST-6×His-SAID1/SAID2* vectors, *SAID1* and *SAID2* were cloned and ligated into EcoRI/KpnI-digested *pAcGHLT-C* (BD Biosciences) by one-step cloning. To generate *pGEX-4T-1-GST-SAID1*_*1-300*_*-6×His* and *pGEX-4T-1-GST-SAID2*_*1-400*_*-6×His* vectors, *SAID1*_*1-300*_ and *SAID2*_*1-400*_ were cloned and ligated with EcoRI/XhoI-digested *pGEX-4T-1-GST-6×His* vectors. Other vectors such as *pET28a-Avi-6×His-SUMO-SE, pET28a-Avi-6×His-SUMO-HYL1, pAcGHLT-GST-6×His-DCL1* etc., were generated in previous work (Wang et al., 2018). Arabidopsis transformations were performed by the floral-dip method, using the ABI or GV3101 Agrobacterium strains.

### Y2H and Y3H assays

The Y2H and Y3H assays were performed as described previously (Wang et al., 2018). The Y2H assays were carried out in the Matchmaker Gold Yeast Two-Hybrid System (Clontech). The pairs of corresponding recombinant constructs fused with *pGBKT7 BD* and *pGADT7 AD* were co-transformed into the Y2H Gold yeast competent cells. The yeast cells co-expressing proteins fused with AD and BD were selected subsequently on the SD-Leu/-Trp double dropout medium, and interaction was examined on SD-Ade/-His/-Leu/-Trp quadruple dropout medium (Clontech, Cat. No. 630317 and 630323). For Y3H assays, the combinations of corresponding *pBridge BD* and *pGADT7 AD* plasmids were co-transformed into the Y2H Gold yeast competent cells. The yeast cells co-expressing proteins fused with pBridge BD and AD were selected subsequently on the SD-Leu/-Trp double dropout medium, and impact of SAID1/SAID2 on SE interaction with other proteins was examined on SD-His/-Leu/-Met/-Trp quadruple dropout medium (Clontech, Cat. No. 630429) supplemented with 5 mM 3-amino-1,2,4-triazole (3-AT, Sigma-Aldrich) and 0 mM, 0.07 mM, 0.14 mM, or 1 mM Met (Sigma-Aldrich). Different amounts of Met (0, 1, and 2 mM) were added to the medium to regulate the transcription activity of Met23 promoter. Note that yeast colonies tend to grow faster in the medium with the nutrient Met than the one without Met. At least 10 individual colonies with similar result were examined for each combination.

### BiFC assay

Protoplasts were prepared from four-week-old leaves and transformed as described (Yoo et al., 2007). The pairs of corresponding recombinant constructs fused with *nYFP* and *cYFP* were co-transformed into protoplasts. Fluorescence signals were visualized 12 hrs after transformation with Leica SP8 inverted confocal microscopy. At least 10 independent protoplasts for each interaction were examined and showed similar results.

### Confocal microscopy

Fluorescence images were taken with Leica SP8 confocal microscope or Olympus Fluoview FV3000 confocal microscope. GFP, YFP, PI, and chlorophyll auto-fluorescence were excited at 488 nm, 514 nm, 561 nm, 640 nm respectively, and the corresponding emission ranges were 500-530 nm, 516-530 nm, 590-620 nm, and 650-700 nm. Sequential module was applied to eliminate potential crosstalk of multiple fluorescent signals.

### Fluorescence recovery after photobleaching

FRAP *in vivo* was performed on Olympus Fluoview FV3000 equipped with a PlanApo N SC2 BFP1 60X oil-immersion objective (Olympus). Nuclear condensates were selected as the region of interest (ROI) and bleached using 0.5 sec pulse of the 488 nm laser for eGFP at full power. The recovery images were taken at equal time intervals of 2 s for 600 s under emission range of 500-530 nm.

### GUS staining

Three-week-old seedlings were sampled and immersed into GUS staining buffer with 2 mM X-Gluc as described previously (Wang et al., 2018). Samples were vacuum-infiltrated for 30 min and incubated at 37 °C for 6hrs. The stained samples were then immersed in 70% ethanol for 2 hrs, and images were taken with an Olympus SZH10 stereo microscope.

### RNA extraction and RT-PCR

Total RNA was extracted from leaves of three-week-old seedlings with TRIzol reagent (Sigma-Aldrich) and treated with TURBO™ DNase (Invitrogen, Cat. No. AM2238). First strand cDNA was reverse transcribed from 1μg of cleaned RNA with SuperScript III reverse transcriptase (Invitrogen, Cat. No. 18080093) with oligo(dT) primers. RT-PCR was performed with primers listed for alternative splicing, with *EF-1α* as internal control. RT-qPCR was performed using iTaq SYBR green Supermix (Bio-Rad) in the Bio-Rad CFX384 Real-Time PCR System. Primers were listed in Supplemental Table 1, with *EF-1α* or *UBQ10* served as internal control. Stem-loop RT-qPCR of miRNA expression was performed as described previously (Zhang et al., 2017). First strand cDNA was synthesized with miRNA/U6-specific RT primers, and miRNA expression was examined with miRNA-specific qPCR forward primers and universal reverse primers, with U6 as internal control. Relative expression level was calculated with the 2^− ΔΔCt^ method (Livak and Schmittgen, 2001).

### sRNA blot assay

Total RNA was extracted from leaves of three-week-old seedlings with TRIzol reagent (Sigma-Aldrich) and treated with TURBO™ DNase (Invitrogen, Cat. No. AM2238). Total RNAs were loaded and sRNAs were separated in 15% denaturing Urea-PAGE gels, followed by semi-dry transferring onto Hybond-N+ hybridization membranes (GE Healthcare, RPN303B). The membranes were UV cross-linked and hybridized with miRNA probes labeled with [γ-^32^P] ATP (PerkinElmer), with U6 served as loading control. Membranes were covered with phosphor imaging plate (GE Healthcare) and signals was detected with Typhoon FLA7000 (GE Healthcare) as described previously (Wang et al., 2018). Probes used for sRNA blot were listed in Supplementary Table 2.

### RNA-seq and sRNA-seq

Total RNA was extracted from leaves of three-week-old seedlings with TRIzol reagent (Sigma-Aldrich) and treated with TURBO™ DNase (Invitrogen, Cat. No. AM2238). For sRNA-seq, total RNA was loaded and sRNA was separated in 15% denaturing Urea-PAGE gels and purified for library generation. For RNA-seq, libraries were generated with TruSeq Stranded Total RNA kit after rRNA depletion with Ribo-Zero Plant (Illumina) based on manufacturer’s guidelines. Library preparation, sequencing, and bioinformatics analysis were performed as described previously (Ma et al., 2018b). Gene ontology analyses were performed on Metascape (Zhou et al., 2019).

### Western blot

Proteins were separated on 8-12% SDS–PAGE gels and transferred onto nitrocellulose membranes (Amersham, 10600003). The primary antibodies used were anti-Actin (Sigma, A0480), anti-Flag (Sigma, F1804) anti-Myc (Sigma, C3956), anti-HA (Sigma, H9658), anti-His (Sigma H1029), anti-GFP (Agrisera, AS15 2987), anti-SE (home made), anti-HYL1(home made), and anti-AGO1(Agrisera, AS09527). The secondary antibodies were ECL Mouse IgG or ECL Rabbit IgG, HRP-linked whole Ab (Amersham, NA931 or NA934). For WB of yeast proteins, ~10^5^ of yeast cells were harvested and resuspended in 200 μL lysis buffer (0.1M NaOH, 0.05 M EDTA, 2 % SDS, 2 % β-mercaptoethanol). The lysates were supplied with 0.1M Acetic Acid and denatured at 95 °C for 10 min. The extracts were then mixed with 50 μL protein loading buffer and subjected for WB.

### Size-exclusion chromatography assay

SEC was performed as previously described(Li et al., 2020c). Three-week-old seedlings were ground to fine powder in liquid nitrogen, and homogenized with three volume of extraction buffer (20 mM Tris-HCl pH 7.5, 300 mM NaCl, 4 mM MgCl_2_, 200 μM ZnCl_2_, 0.1% Triton X-100, 1% glycerol, 4X cOmplete™ EDTA-free protease inhibitor, 2 mM PMSF and 15 μM MG132).

The homogenates were centrifuged twice at 15,000 rpm for 15 min at 4 °C. The supernatant was then filtered through 0.2 μm filter and loaded into the Superdex 200 10/300 GL column (GE Healthcare) prewashed with balance buffer (20 mM Tris-HCl pH 7.5, 300 mM NaCl, 4 mM MgCl_2_, 200 μM ZnCl_2_, 0.1% Triton X-100, 1% glycerol, 1/3 X cOmplete™ EDTA-free protease inhibitor, 0.5 mM PMSF and 15 μM MG132). The running buffer contained 20 mM Tris-HCl pH 7.5, 300 mM NaCl, 4 mM MgCl_2_, 200 μM ZnCl_2_, 0.1% Triton X-100, 1% glycerol, 1X cOmplete™ EDTA-free protease inhibitor, 2 mM PMSF and 15 μM MG132. Fractions of 0.6 mL were collected for western blot analysis using an anti-SE antibody for SE. The Superdex 200 10/300 GL column was calibrated with the gel filtration protein standard (Bio-Rad).

### In vitro cell-free decay assay

Three-week-old seedlings were ground to fine powder in liquid nitrogen, and homogenates were prepared of 0.4g of the powder in 1.6mL of protein lysis buffer (25 mM Tris-HCl pH 7.5, 10 mM NaCl, 10 mM MgCl_2_ and 10% glycerol). The homogenates were centrifuged twice at 4 °C for 10 min at 15,000 rpm and adjusted to equal volume with the protein lysis buffer. The lysates were treated with 0.5 mM CHX and aliquot into two parts. One aliquot was treated with 50 µM MG-132 (Calbiochem, Cat. NO. 474787), and the other aliquot was treated with 2% DMSO as control. The mixtures were then incubated at room temperature for the indicated time points (0, 10, 20, 30, 45, 60, 75 min), and collected for WB with an anti-SE antibody, as described previously (Wang et al., 2022).

### Phos-tag SDS-PAGE assay

Three-week-old seedlings were ground to fine powder in liquid nitrogen. The lysates were prepared with 0.1 g powder homogenized with 1 mL of 10% (m/V) TCA/acetone by precipitation at 4 °C for 30 min. The lysates were subjected for Phos-tag™ SDS-PAGE analysis following guidelines of the manufacture (Nard Institute, LTD, Phos-tag™ Acrylamide AAL-107M).

### In vitro kinase assay

The in vitro kinase assays were performed as previously described (Wang et al., 2022). Briefly, 25 nM kinase, 350 nM recombinant SE, and SAID1/2 or their truncated variants with indicated concentrations were incubated in 30 μL reaction buffer (10 mM Tris-HCl, pH7.5, 5 mM MgCl_2_, 2.5 mM EDTA, 50 mM NaCl, 0.5 mM DTT, 50 μM ATP and 1 μCi [γ-^32^P]ATP) at room temperature for indicated time duration or 1hr. Samples were denatured with 4 X SDS loading buffer and separated by 8% SDS-PAGE gel. Phosphorylation was analyzed by autoradiography.

### Expression and purification of recombinant proteins

Protein expression and purification were performed as described previously (Wang et al., 2022; Wang et al., 2018). The recombinant proteins 6×His-SUMO-SE, 6×His-SUMO-HYL1, GST-SAID1_1-300_-6×His, and GST-SAID2_1-400_-6×His were expressed in E. coli BL21 (DE3) cells. Briefly, for purification of recombinant proteins in *E. coli* system, cells were induced with 0.5 mM Isopropyl-β-D-thiogalactopyranoside (IPTG) for 3 hours at 37°C when OD_600_ of cultured BL21 cells reached 0.6. Pellets were collected and re-suspended in 50mL lysis buffers with minor modification for each protein (For SE: 20 mM Tris-HCl pH 8.5, 500 mM KCl, 1 mM β-mercaptoethanol, 2 mM PMSF, 2% glycerol, 0.1% Triton X-100, 1 pellet per 50 mL cOmplete™ EDTA-free protease inhibitor. For HYL1: 40 mM Tris-HCl pH 8.0, 300 mM KCl, 1 mM β-mercaptoethanol, 1 mM PMSF, 2% glycerol, 1% Triton X-100, 1 pellet per 50 mL cOmplete™ EDTA-free protease inhibitor. For SAID1_1-300_-6×His and GST-SAID2_1-400_-6×His: 40 mM Tris-HCl pH 8.0, 300 mM KCl, 1 mM β-mercaptoethanol, 2 mM PMSF, 2% glycerol, 0.1% Triton X-100, 1 pellet per 50 mL cOmplete™ EDTA-free protease inhibitor). The cells were lysed by French Press at 25,000 psi (Avestin, EmulsiFlex-C3) and then centrifuged at 18,000 rpm for 15 min at 4 °C. The supernatant was filtrated through 0.45 µm filter and loaded into HisTrap HP column. The collected fractions were concentrated by Amicon® Ultra-15 Centrifugal Filter Units (EMD Millipore) and subjected for gel filtration by HiLoad^®^ 16/600 Superdex^®^ 200 pg column (GE Healthcare). The recombinant proteins were dialyzed in dialysis buffer (20 mM Tris-HCl pH 8.5, 60 mM KCl, 50% glycerol, 2 mM DTT, 2 mM β-mercaptoethanol) at 4 °C overnight, and then aliquot and stored at −80 °C.

GST-6×His-PRP4KA, GST-6xHis-DCL1, GST-6xHis-SAID1, and GST-6xHis-SAID2 recombinant proteins were expressed and induced in sf9 insect cell using BD BaculoGold™ Baculovirus Expression System. (BD Biosciences, 554738 and 554740). The corresponding *pAcGHLT-C-GST-6xHis* plasmids and BaculoGold baculovirus DNA were co-transfected into sf9 insect cells to generate the recombinant baculovirus. The recombinant viruses were amplified and the resultant cells containing P3 virus was inoculated into 2.5×10^6^ sf9 insect to propagate until proteins were sufficiently expressed. The pellets were collected and re-suspended in lysis buffer buffers with minor modification for each protein (For DCL1: 100 mM Tris-HCl pH 8.0, 300 mM KCl, 2% glycerol, 1 mM β-mercaptoethanol, 1 mM PMSF, 1% Triton X-100, 1 pellet per 50 mL cOmplete™ EDTA-free protease inhibitor. For PRP4KA: 40 mM Tris-HCl pH 7.5, 500 mM NaCl, 5% glycerol, 1 mM β-mercaptoethanol, 1 mM PMSF, 0.1% Triton X-100, 1 pellet per 50 mL cOmplete™ EDTA-free protease inhibitor. For SAID1 and SAID2: 40 mM Tris-HCl pH 8.5, 300 mM KCl, 2% glycerol, 0.1% Triton X-100, 1 mM β-mercaptoethanol, 50μM MG132, 3 mM PMSF, 1 pellet per 50 mL cOmplete™ EDTA-free protease inhibitor). The cells were lysed by French Press at 25,000 psi (Avestin, EmulsiFlex-C3) and then centrifuged at 18,000 rpm for 15 min at 4 °C. The supernatant was filtrated through 0.45 µm filter and loaded into HisTrap HP column (GE Healthcare, Cat#:17-5248-02). The collected fractions were concentrated by Amicon® Ultra-15 Centrifugal Filter Units (EMD Millipore) and subjected for gel filtration by HiLoad^®^ 16/600 Superdex^®^ 200 pg column (GE Healthcare). Correct peaks were collected and the recombinant proteins were dialyzed in dialysis buffer (20 mM Tris-HCl pH 8.5, 60 mM KCl, 50% glycerol, 2 mM DTT, 2 mM β-mercaptoethanol) at 4 °C overnight, and then aliquot and stored at −80 °C.

### In vitro transcription and 5′ labelling of RNAs

In vitro transcription and 5′ labelling of RNA substrates were performed as described (Zhu et al., 2013). RNA substrates were transcribed under the T7 promoter *in vitro* using PCR-generated pri-miRNA templates or the ssRNA G3A44 template. Primers used for PCR of template and in vitro transcription of RNA substrates are listed in Supplementary Table 2. For 5′-end labeling of RNAs, 5 pmol of label-free RNA substrates were dephosphorylated with Alkaline Phosphatase (New England Biolabs) and labeled with [γ-^32^P] ATP (PerkinElmer) by T4 Polynucleotide Kinase (New England Biolabs). The labeled RNAs were fractionated with 6% denaturing Urea-PAGE gels and extracted from gel slices. The labeled RNAs were heated at 95 °C for 3 min and naturally cooled down to room temperature for RNA annealing. Final concentrations were adjusted to 1000 cpm μL^−1^ for each RNA substrate.

### Electrophoretic mobility shift assay

EMSA assays were performed as described previously (Wang et al., 2018). Recombinant proteins and 5′-end labeled RNAs were incubated in the EMSA buffer (20 mM Tris-HCl pH 7.5, 2 mM MgCl_2_, 2 mM DTT, 5 mM ATP, 0.3% NP-40, 1 U/μL SUPERase-In RNase Inhibitor) for 30 min at room temperature. The final concentration of NaCl or KCl was ~55 mM from protein/RNA dialysis buffer. The RNA–protein complexes were resolved on native 1% agarose gels and visualized by radiography after RNA fixation in RNA fixation buffer (40% ethanol, 10% acetic acid, and 5% glycerol) for 20 min and dried at 80 °C for 2 hrs. The Kd and appKd were calculated using GraphPad Prism 7 software fit with a Hill slope model.

### In vitro microprocessor assay

In vitro dicing activity assays were performed as described previously (Wang et al., 2018). Briefly, 1,000 cpm of RNA substrates, 0.1 pmol SE, 0.2 pmol HYL1, 0.05 pmol DCL1, various concentrations of SAID1/SAID2 recombinant proteins or their truncated variants were incubated in 30 μL assay buffer containing 20 mM Tris-HCl pH7.5, 50 mM KCl, 4 mM MgCl_2_, 1 mM DTT, 5 mM ATP, 1 mM GTP and 1 U/μL SUPERase-In RNase Inhibitor (Thermo Fisher). The final concentration of NaCl and KCl was ~70 mM, with ~20 mM salt from the protein/RNA dissolving buffer. The dicing reactions were performed at 37 °C for 30 min and stopped by adding 1 volume TBE-Urea sample buffer (Bio-Rad), heating at 95 °C for 10 min and quickly chilling on ice for 2 min. The processed mixtures were fractionated on 10% denaturing Urea-polyacrylamide gels. Gels were dried at 80 °C for 2 hrs and covered with phosphor imaging plate (GE Healthcare) overnight after RNA fixation in RNA fixation buffer (40% ethanol, 10% acetic acid, and 5% glycerol) for 20 min. Signals were detected and captured with Typhoon FLA7000 (GE Healthcare). Quantification of the Microprocessor cleavage efficiency was calculated by the ratio of processed to unprocessed pri-miRNA fragments. The relative efficiency was normalized to that of the Microprocessor reaction control whose ratio was arbitrarily set at 1. Data are means ± SD of three replicates. Plots and fitting by exponential equation or linear regression were performed with GraphPad Prism 7.

### Ribonucleoprotein immunoprecipitation (RIP) assay

RIP assay was performed as previously described (Wang et al., 2018) with GFP-Trap Agarose beads (ChromoTek, Cat no. gta). Briefly, ten-day-old Col-0, *pSAID1::GFP:SAID1*, and *pSAID2::GFP:gSAID2* seedlings were cross-linked with 0.1% formaldehyde and ground to fine powder in liquid nitrogen. Nuclei were isolated and homogenized with RIP buffer (50 mM Tris-HCl pH 8.5, 150 mM KCl, 5 mM MgCl_2_, 0.5% Triton X-100, 2% glycerol, 0.25% NP-40, 1% SDS, 5 mM DTT, 1 mM PMSF, 50 µM MG-132, 1 pellet per 10 mL Complete EDTA-free protease inhibitor (Roche) with 10 U/ml TURBO DNase (Thermo Fisher)). After centrifugation at 15,000 RPM for 15 min, the supernatant was immunoprecipitated with GFP-Trap Agarose beads or Sepharose beads without antibody control at 4 °C for 2 hrs. Immunoprecipitates were washed three times with RIP buffer and once with high salt buffer (50 mM Tris-HCl pH 8.5, 500 mM KCl, 5 mM MgCl_2_, 5 mM DTT, 0.2% Triton X-100, 2% glycerol, 1 mM PMSF, 50 µM MG-132, 1 pellet per 10 ml Complete EDTA-free protease inhibitor (Roche)) at 4 °C for 5 min. After treatment with Proteinase K, the RNA was extracted using TRI Reagent (Sigma-Aldrich) and treated with TURBO DNase (Thermo Fisher). RNA was reverse-transcribed and used for RT–qPCR of pri-miRNAs. The relative enrichment of pri-miRNAs was normalized to the control without antibody after normalization with the input, where the ratio was arbitrarily set to 1 with s.d. from three biological repeats (\**P* < 0.05; \*\**P* < 0.01; unpaired, two-tailed Student’s *t-*test).

### Quantification and statistical analysis

For RNA-seq and sRNA-seq, DESeq2 package was used to normalize gene expression levels with trimmed mean of M-values according to the false discovery rate (FDR), with the cutoff of significance as 0.05. The band intensity of Western blot, sRNA blot, EMSA, and microprocessor activity were quantified with ImageJ. Curves were fitted and values were calculated with GraphPad Prism 7 in EMSA and Microprocessor activity assays. Unpaired two-tailed Student’s *t* test was performed to calculate the P value. The cutoff for significance was 0.05. *P < 0.05, **P < 0.01, and ***P < 0.001.

### Model diagram

The model diagram was created on Biorender.

